# Butyrate supplementation reshapes systemic metabolism and attenuates cardiac aging

**DOI:** 10.64898/2026.06.08.730600

**Authors:** Mingming Tong, Lihua Jiang, Ruiqi Jian, Faye Chleilat, Frank Wong, Tristan Chou, Lettie McGuire, Michael P. Snyder

## Abstract

Identifying interventions that improve health and cardiac function is important for extending healthspan. We show that supplementation with the SCFA butyrate beginning at 18 months (late life) extends survival in male and female mice and improves systemic metabolic and physical-performance measures. Treatment reduces adiposity and fasting glucose and attenuates age-associated cardiac hypertrophy, fibrosis, and systolic and diastolic dysfunction. Integrated plasma and myocardial metabolomics and lipidomics reveal compartment-dependent metabolic remodeling, while cardiac proteomics and full-length RNA sequencing identify coordinated changes in extracellular-matrix, RNA-processing, lipid-metabolism, and autophagy-associated programs. These molecular responses are accompanied by improved mitochondrial ultrastructural organization and respiratory function. Butyrate supplementation also enhanced cardiac autophagy and mitophagy, and its beneficial effects on LV hypertrophy and diastolic indices required ATG7. These findings demonstrate that butyrate supplementation initiated at an advanced age modifies systemic and cardiac aging phenotypes and implicate metabolic remodeling and mitochondrial quality control pathways as contributing to these benefits.

**Graphical abstract:** 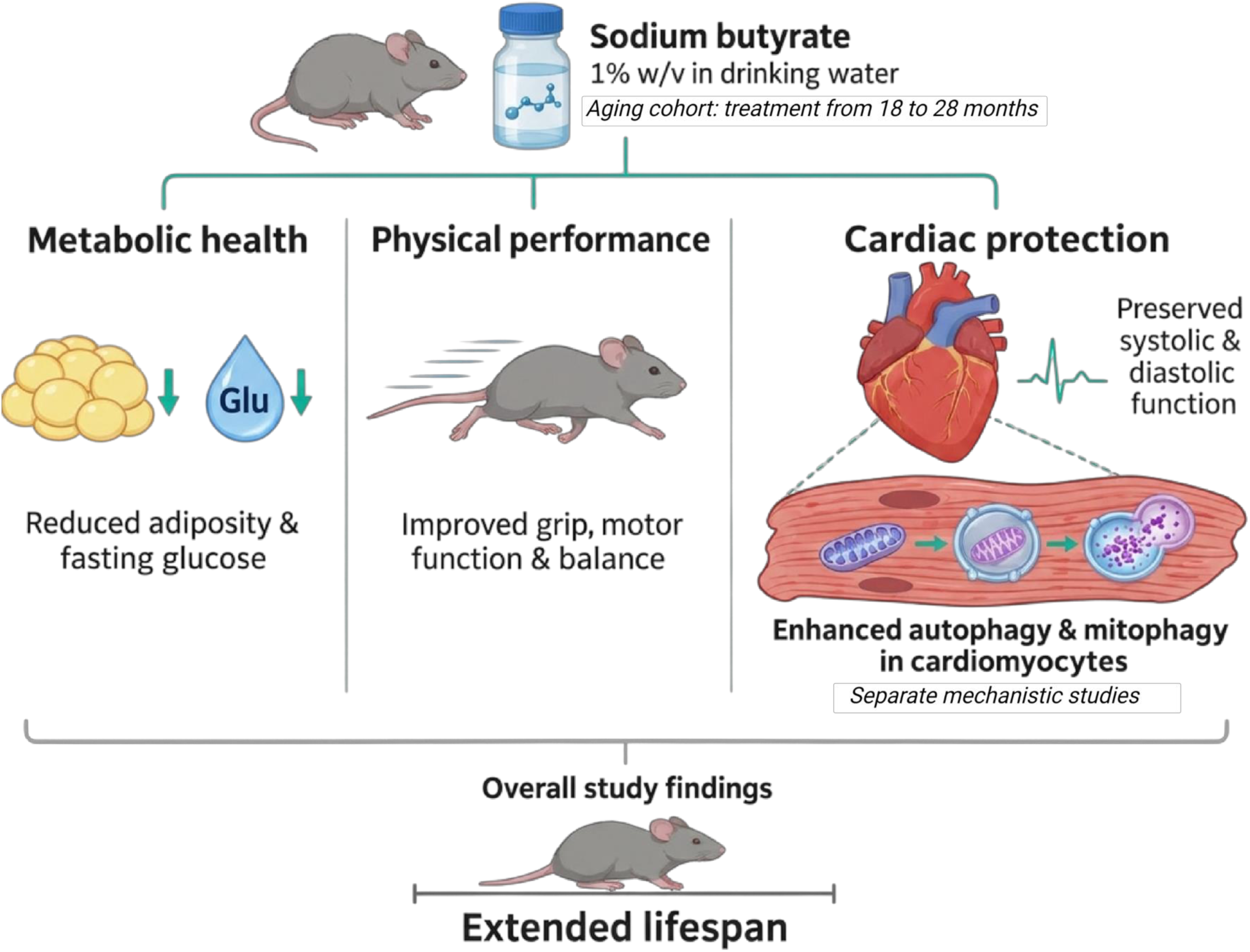

## Introduction

Aging is accompanied by progressive changes in metabolic homeostasis, tissue maintenance, and organ function that increase vulnerability to chronic disease and mortality^1, 2^. The cardiovascular system in particular undergoes age-associated structural and molecular remodeling, including myocardial hypertrophy, extracellular matrix (ECM) accumulation, mitochondrial dysfunction, and alterations in cellular quality control pathways^3–6^. These interconnected processes contribute to impaired ventricular relaxation, reduced cardiac reserve, and increased susceptibility to cardiovascular disease, sometimes before substantial deterioration in systolic performance^7, 8^. Although interventions targeting fundamental aging processes have attracted increasing interest, whether metabolic interventions initiated after substantial age-associated changes have accumulated can attenuate cardiac aging remains unknown. Defining the capacity of the aging heart to respond to interventions introduced late in life may reveal opportunities to preserve cardiovascular function and improve physiological resilience.

Diet and the gut microbiota are important determinants of host metabolism and molecular regulation. Short-chain fatty acids (SCFAs), including acetate, propionate, and butyrate, are produced primarily through microbial fermentation of dietary substrates and participate in communication between the microbiota and host^9–11^. Our previous studies demonstrated that dietary fiber interventions elicit individualized microbial and host molecular responses in human and mice^12^ and that butyrate and propionate can influence chromatin-associated regulation and gene expression through epigenetic mechanisms^13^. Extending this work to cardiovascular disease, our recent integrated gut microbiome and host multi-omics study in patients with chronic systolic heart failure identified microbial and metabolic signatures associated with disease severity and clinical improvement^14^. Together, these findings establish a connection between dietary factors, microbial metabolites, host molecular regulation, and cardiovascular physiology. However, whether supplementation with an individual microbiota-associated metabolite can modify established features of cardiac aging, particularly when initiated late in life, remains unclear.

Butyrate is of particular interest because its biological effects extend beyond its role as an energy substrate. Experimental studies have demonstrated effects on glucose and energy metabolism, substrate utilization, immune regulation, and gene expression, including mechanisms involving histone deacetylase inhibition^13, 15, 16^. Previous studies have also reported that sodium butyrate attenuates cardiac remodeling and dysfunction in experimental cardiometabolic disease^17^, while emerging clinical trials suggest that oral supplementation can influence adiposity and glucose regulation in selected populations^18^. Nevertheless, whether these effects extend to naturally aged hearts and whether prolonged supplementation initiated at an advanced age can modify cardiac structural and functional deterioration remain unresolved. The heart has high energetic demands and specialized requirements for substrate utilization, mitochondrial metabolism, and lipid homeostasis, all of which undergo substantial remodeling during aging^3, 11, 19, 20^. Moreover, circulating metabolic changes may not directly reflect myocardial responses, highlighting the importance of examining systemic and cardiac metabolism together. Integrating plasma and myocardial metabolomics and lipidomics can therefore help distinguish systemic metabolic responses from changes occurring within cardiac tissue. In parallel, age-associated cardiac remodeling involves changes in ECM organization, RNA processing, mitochondrial integrity, and autophagy-and mitophagy-associated pathways^3–6, 21–27^. Because autophagy and mitophagy contribute to the removal of damaged intracellular components and maintenance of mitochondrial homeostasis, their dysregulation may connect metabolic stress to structural and functional cardiac decline^21–23^. Integrating proteomics with full-length RNA sequencing provides additional opportunities to identify molecular changes that are not fully captured by gene-level expression measurements^24, 28, 29^. However, whether late-life butyrate supplementation coordinately modifies these systemic, myocardial, and intracellular quality-control processes remains unknown.

Here, we investigated the effects of sodium butyrate supplementation initiated at 18 months of age in male and female C57BL/6J mice. We performed survival and systemic physiological assessments along with cardiac phenotyping, plasma and left ventricular (LV) metabolomics and lipidomics, cardiac proteomics, PacBio Kinnex full-length RNA sequencing, mitochondrial ultrastructural analysis, and respiratory measurements in isolated cardiac mitochondria. We further assessed autophagy- and mitophagy-associated activity in separate age-defined cohorts and examined the contribution of ATG7-dependent processes to selected cardiac responses in a specific model of heart failure with preserved ejection fraction (HFpEF). By integrating organism-level, tissue-level, and molecular measurements, we sought to determine whether late-life butyrate supplementation modifies survival, physiology and cardiac aging phenotypes, and we characterize compartment-dependent metabolic remodeling, and identify candidate pathways connecting metabolic regulation, mitochondrial maintenance, and cardiac function.

## Results

### Late-life butyrate supplementation extends survival and improves metabolic and physical function in aged mice

To determine whether metabolic intervention initiated late in life could attenuate age-associated physiological decline, we administered 1% sodium butyrate in drinking water to male and female C57BL/6J mice beginning at 18 months of age. The experimental design integrated longitudinal systemic phenotyping, cardiac functional assessments, plasma and myocardial multi-omics profiling, and mitochondrial analyses, together with independent survival cohorts and complementary mechanistic studies (Figure 1A). Given the relatively long lifespan of this strain and the progressive deterioration of metabolic and physical function during aging, we assessed survival, systemic butyrate exposure, adiposity, glucose homeostasis, respiratory metabolism, and physical performance as complementary measures of late-life physiological health^30–33^.

**Figure 1.**
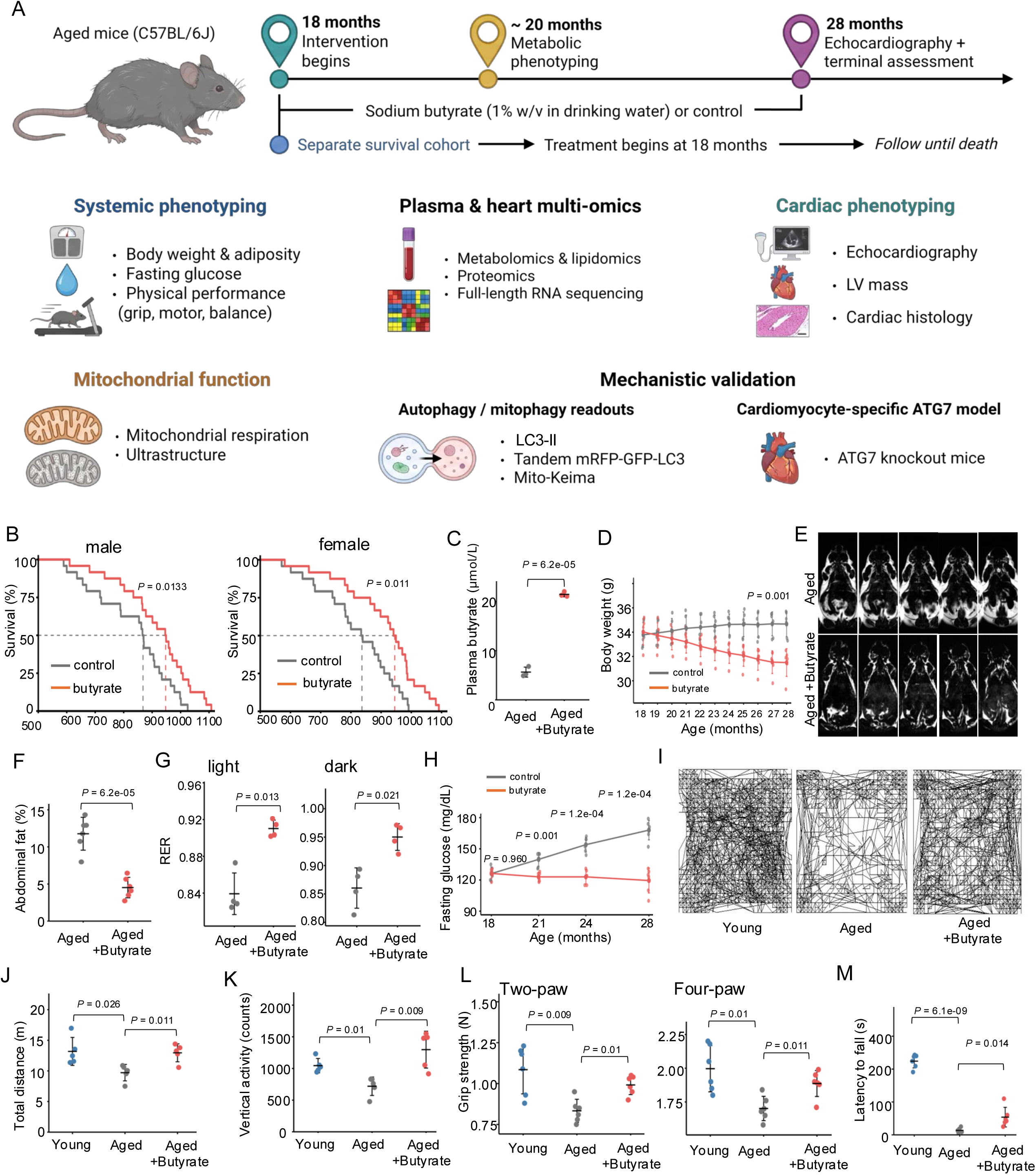
Late-life sodium butyrate supplementation extends male and female survival and improves systemic metabolic and physical phenotypes during aging. (A) Schematic overview of the experimental design and study workflow. Male and female C57BL/6J mice received sodium butyrate (1% w/v) in drinking water or control water beginning at 18 months of age. Longitudinal physiological assessments were performed during treatment, with cardiac phenotyping and terminal tissue analyses conducted at 28 months. Major assessments included systemic metabolic and physical phenotyping, plasma and myocardial multi-omics profiling, cardiac structure and function, and mitochondrial ultrastructural and respiratory analyses. Independent survival cohorts were followed until death. Additional autophagy and mitophagy reporter experiments and cardiomyocyte-specific ATG7-deficient mouse studies were performed in separate cohorts to investigate potential mechanisms. (B) Kaplan–Meier survival curves for male and female control and sodium butyrate-treated mice (n = 24 per group per sex). Dashed lines indicate the 50% survival threshold and estimated median survival times. P values were determined using the log-rank test. (C) Plasma butyrate concentrations in aged male control and butyrate-treated mice, measured using targeted LC–MS/MS (n = 3 per group). (D) Longitudinal body weight trajectories in male control and butyrate-treated mice from 18 to 28 months of age (n = 6 per group). The indicated P value represents the between-group comparison at 28 months. (E) Representative coronal anatomical magnetic resonance images showing five consecutive slices from one aged control mouse and one butyrate-treated aged male mouse at 28 months of age. Images are presented for qualitative anatomical comparison and were not used for quantitative adiposity measurements. (F) Abdominal fat index, calculated as the combined weight of dissected epididymal and perirenal adipose tissue divided by total body weight and expressed as a percentage (n = 6 per group). (G) Respiratory exchange ratio (RER; VCO₂/VO₂) during the light and dark phases in aged male control and butyrate-treated mice (n = 4 per group). (H) Longitudinal fasting blood glucose concentrations measured at 18, 21, 24, and 28 months of age in male control and butyrate-treated mice (n = 6 per group). Indicated P values represent age-specific between-group comparisons. (I) Representative open-field movement trajectories recorded over 60 min in young, aged control, and aged butyrate-treated male mice. (J and K) Total distance traveled (J) and vertical activity counts (K) during open-field testing in young, aged control, and aged butyrate-treated male mice (n = 5 per group). (L) Two-paw and four-paw grip strength in young, aged control, and aged butyrate-treated male mice (n = 6 per group). Both measurements were obtained from the same animals. (M) Latency to fall during the final trial of an accelerating rotarod protocol in young, aged control, and aged butyrate-treated male mice (n = 6 per group). Young reference mice were 4 months old, and aged mice were 28 months old at physical-performance assessment. The indirect-calorimetry cohort was approximately 20 months old. Individual data points represent biologically independent mice, and summary marks and error bars indicate mean ± SD. Longitudinal plots connect group means. Survival distributions were compared using the log-rank test. Other endpoints were analyzed using two-sided, independent, equal-variance Student’s t-tests, with BH correction.

In independent survival cohorts, late-life butyrate supplementation significantly extended survival in both male (P = 0.0133) and female mice (P = 0.011), with 24 mice per group for each sex (Figure 1B). Median survival increased by approximately 10% in treated males and females relative to their respective controls. Targeted plasma measurements confirmed substantially higher circulating butyrate concentrations in treated aged males (21.49 μmol/L) than in untreated aged controls (5.65 μmol/L; P = 6.2 × 10^-5^; Figure 1C). Thus, supplementation initiated at an advanced age increased systemic butyrate exposure and was associated with prolonged survival in both sexes.

We next investigated whether the survival benefit was accompanied by improvements in systemic metabolic health. Longitudinal body-weight monitoring revealed divergent trajectories in male mice, with body weight remaining relatively stable or increasing modestly in controls but declining progressively in butyrate-treated animals (Figure 1D). At 28 months, treated males weighed significantly less than aged controls (31.50 versus 34.67 g; P = 0.001).

Representative whole-body magnetic resonance images (MRI) from untreated aged and butyrate-treated aged mice are shown as five consecutive anatomical slices from one mouse per group (Figure 1E), providing a qualitative comparison of body fat distribution. Consistent with the body-weight differences, dissected abdominal adipose tissue measurements revealed a marked reduction in adiposity following butyrate supplementation. The abdominal fat index, calculated as the combined mass of epididymal and perirenal adipose tissue normalized to total body weight, was significantly lower in treated aged males (P = 6.2 × 10^-5^; Figure 1F), accompanied by a reduction in absolute abdominal fat mass (Supplementary Figure 2A). Female mice exhibited a similar reduction in body weight, with mean values of 28.42 g in treated mice compared with 32.83 g in controls at 28 months (P = 6.0 × 10^-4^; Supplementary Figure 1A). Together, these observations indicate that late-life butyrate supplementation was associated with reduced abdominal fat accumulation and altered body-weight trajectories during aging.

To characterize whole-body metabolic responses, we performed indirect calorimetry in a separate cohort of approximately 20-month-old male mice. Body weight during metabolic monitoring did not differ significantly between groups (Supplementary Figure 2B), nor did daily water intake or light- and dark-phase food intake (Supplementary Figure 2C and D). In contrast, butyrate-treated mice exhibited a significantly higher respiratory exchange ratio (RER) during both the light (P = 0.013) and dark (P = 0.021) phases (Figure 1G), consistent with an improved shift in relative metabolic substrate utilization toward carbohydrate oxidation under the measured conditions. Carbon dioxide production (VCO₂) was significantly higher during the dark phase (P = 0.016), whereas oxygen consumption (VO₂) showed a trend toward higher values without reaching statistical significance (Supplementary Figure 2E and F, P=0.058). Body-weight-normalized heat production was also higher during the dark phase (P = 0.041; Supplementary Figure 2G). These findings indicate that butyrate supplementation altered whole-body respiratory metabolism.

Changes in body composition and respiratory metabolism were accompanied by lower fasting glucose concentrations. In male controls, fasting blood glucose increased progressively from 18 to 28 months of age, whereas concentrations remained comparatively stable in butyrate-treated mice (Figure 1H). At 28 months, fasting glucose was significantly lower in treated males than in controls (119.5 versus 168.0 mg/dL; P = 1.2 × 10^-4^). Female mice exhibited a similar pattern, with concentrations of 127.50 mg/dL in treated mice compared with 160.67 mg/dL in controls at 28 months (P = 9.5 × 10^-5^; Supplementary Figure 1B). These results suggest that supplementation attenuated the age-associated increase in fasting glucose in both sexes.

We next examined whether these metabolic differences were accompanied by changes in physical performance. Compared with young reference mice, aged control males exhibited lower spontaneous locomotor activity, including reduced total distance traveled and vertical activity during open-field testing (Figure 1I–K). Butyrate-treated aged males displayed significantly greater total distance traveled (P = 0.011) and vertical activity (P = 0.009) than aged controls (Figure 1J and K). Both two-paw and four-paw grip strength were also higher in treated aged males than in aged controls (P = 0.010 and P = 0.011, respectively; Figure 1L). Motor performance assessed using an accelerating rotarod was similarly improved, with mean latency to fall increasing from 13.5 s in aged controls to 53.8 s in treated aged males (P = 0.014), compared with 224 s in young reference mice (Figure 1M). Female mice exhibited corresponding improvements in two-paw and four-paw grip strength (P = 7.6 × 10^-4^ and P = 0.002, respectively) and rotarod performance (P = 0.004; Supplementary Figure 1C and D). Although performance remained below young-reference levels, these findings demonstrate improvements across multiple measures of physical function following supplementation. Collectively, these results demonstrate that sodium butyrate supplementation initiated at 18 months of age extends survival in male and female C57BL/6J mice and is associated with reduced abdominal adiposity, lower fasting glucose, altered respiratory metabolism, and improved physical performance during advanced aging.

### Late-life butyrate supplementation selectively remodels the circulating metabolome and lipidome

To characterize systemic metabolic changes associated with late-life butyrate supplementation, we performed targeted organic acid profiling, broad metabolomic profiling, and lipidomic analysis of plasma from young (4-month-old), aged control (28-month-old), and aged butyrate-treated (28-month-old) male mice. Given the roles of circulating metabolites in interorgan metabolic communication, these analyses were designed to identify age-associated metabolic signatures and determine their responses to butyrate supplementation^11, 19, 20^.

Targeted profiling using 3-nitrophenylhydrazine (3-NPH) derivatization coupled with liquid chromatography–tandem mass spectrometry revealed age- and treatment-associated differences in central-carbon metabolites, SCFAs, hydroxybutyrates, and other organic acids (Figure 2A). Several tricarboxylic acid (TCA) cycle intermediates, including citrate, α-ketoglutarate, and succinate, were more abundant in aged control plasma than in young plasma but showed lower abundances following butyrate supplementation (Figure 2B). The treatment-associated decrease in α-ketoglutarate brought its abundance closer to young-reference levels, whereas citrate and succinate remained elevated relative to young mice. Similar patterns were observed for acetate, 2-hydroxybutyrate, and 3-hydroxybutyrate, which increased with aging and decreased following supplementation. These findings indicate selective remodeling of circulating organic acid profiles rather than uniform restoration to young-reference abundances.

**Figure 2.**
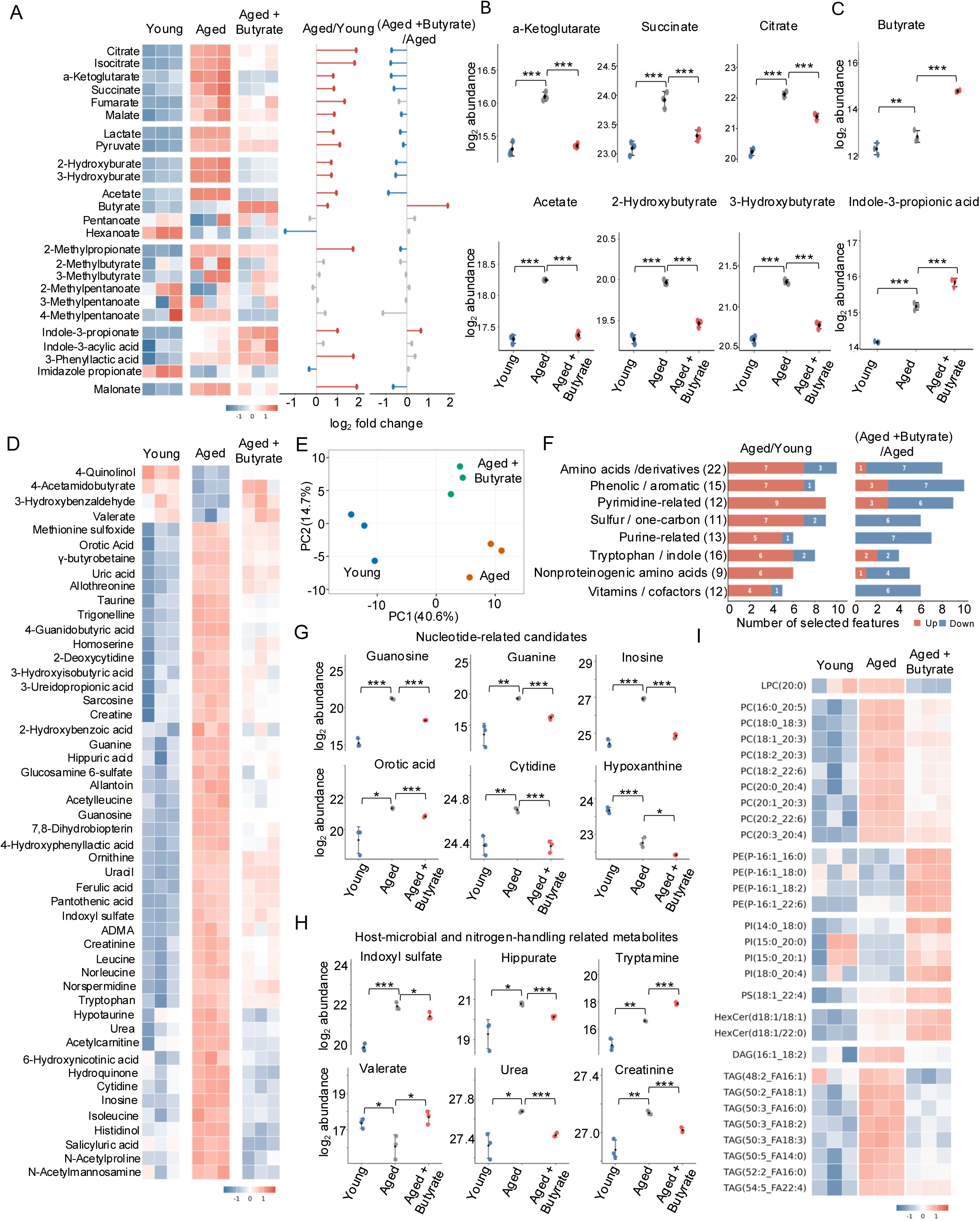
Late-life sodium butyrate supplementation selectively remodels the circulating metabolome and lipidome in aged male mice. Plasma metabolomic and lipidomic analyses were performed in young (4-month-old), aged control (28-month-old), and aged sodium butyrate-treated (28-month-old) male C57BL/6J mice (n = 3 biologically independent mice per group for each analytical platform). (A) Heatmap of selected central-carbon metabolites, short-chain fatty acids, hydroxybutyrates, and related organic acids measured by targeted 3-nitrophenylhydrazine (3-NPH) derivatization LC–MS/MS. Rows represent metabolites and columns represent individual mice. Heatmap colors indicate row-standardized abundance. Adjacent plots show log₂ fold changes for aged control versus young and aged butyrate-treated versus aged control mice. Positive and negative values indicate increased and decreased abundance, respectively, in the numerator group of each comparison. (B) Representative plasma metabolites identified by targeted organic acid profiling, including α-ketoglutarate, succinate, citrate, acetate, 2-hydroxybutyrate, and 3-hydroxybutyrate. Abundances are displayed on a log₂ scale. (C) Plasma butyrate and indole-3-propionic acid (IPA) abundances measured using targeted organic acid profiling. Abundances are displayed on a log₂ scale. (D) Heatmap of selected plasma metabolites identified through broader metabolomic profiling, including amino acid derivatives, organic acids, nucleotide-related compounds, aromatic metabolites, and host–microbial co-metabolites. Rows represent metabolites and columns represent individual mice; colors indicate relative abundance standardized within each metabolite. (E) Principal component analysis (PCA) of plasma metabolomic profiles from young, aged control, and aged butyrate-treated mice. Each point represents an individual mouse. Principal components 1 and 2 explain 40.6% and 14.7% of total variance, respectively. (F) Distribution of significantly altered plasma metabolites across biochemical categories in aged control versus young and aged butyrate-treated versus aged control mice. Red and blue bars indicate the numbers of significantly increased and decreased metabolites, respectively, within each category. Numbers within bars denote metabolite counts. Statistical significance was determined using two-sided Student’s t-tests with Benjamini–Hochberg correction for multiple comparisons (adjusted P < 0.05). (G) Representative purine-and pyrimidine-associated metabolites, including guanosine, guanine, inosine, orotic acid, cytidine, and hypoxanthine. (H) Selected host–microbial co-metabolites and nitrogen-handling-associated compounds, including indoxyl sulfate, hippurate, tryptamine, valerate, urea, and creatinine. (I) Heatmap of selected circulating lipid species spanning lysophosphatidylcholines (LPCs), phosphatidylcholines (PCs), phosphatidylethanolamines (PEs), phosphatidylinositols (PIs), phosphatidylserines (PSs), hexosylceramides (HexCers), diacylglycerols (DAGs), and triacylglycerols (TAGs). Rows indicate lipid species and columns represent individual mice. Colors indicate row-standardized relative abundance. Individual points represent biologically independent mice, and data are presented as mean ± SD where applicable. Differential abundance was evaluated using two-sided, independent, equal-variance Student’s t-tests on log₂-transformed abundance values. BH correction was performed separately across testable features within each assay, sex, and contrast. Volcano plot colors denote nominal P < 0.05, and black outlines identify features additionally meeting BH-adjusted FDR < 0.05. Heatmap colors represent row-standardized abundance values. * *P* < 0.05; ** *P* < 0.01; *** *P* <0.001.

As expected, circulating butyrate was significantly increased in treated aged mice (Figure 2C), consistent with independently quantified plasma butyrate concentrations shown in Figure 1B. However, this increase was not accompanied by a generalized elevation of other SCFAs. Acetate was more abundant in aged controls than in young mice and decreased following treatment, whereas other straight-chain and branched-chain fatty acids exhibited distinct responses (Figure 2A and B). These observations indicate that supplementation with an individual SCFA was accompanied by broader but metabolite-specific changes in circulating organic acid profiles. Such metabolite-specific responses are consistent with the diverse contributions of SCFAs to host energy metabolism and metabolic regulation^9-11, 34^.

Among the affected metabolites, 2-hydroxybutyrate and 3-hydroxybutyrate showed particularly pronounced differences between aged control and butyrate-treated mice (Figure 2B). Although structurally related, these metabolites reflect distinct metabolic processes. 2-Hydroxybutyrate is associated with amino acid metabolism and redox-related metabolic responses^35^, whereas 3-hydroxybutyrate (β-hydroxybutyrate) is a circulating ketone body that can serve as both an energy substrate and a signaling metabolite^36^. Higher circulating 3-hydroxybutyrate concentrations have also been associated with increased mortality in patients with chronic heart failure with reduced ejection fraction^37^. In the present study, the reduction in both hydroxybutyrates following butyrate supplementation suggests changes in systemic intermediary metabolism.

In contrast, indole-3-propionic acid (IPA), a gut microbiota-derived tryptophan metabolite, was elevated in aged controls relative to young mice and increased further following butyrate supplementation (Figure 2C). IPA has been implicated in improved cardiovascular homeostasis in experimental studies. IPA administration was reported to attenuate diastolic dysfunction and cardiac metabolic remodeling in a mouse model of HFpEF^38^, and a separate study identified an association between IPA-mediated regulation of macrophage autophagy and reduced aging-related myocardial fibrosis^39^. In addition, integrated gut microbiome and metabolomic profiling in patients with chronic systolic heart failure associated IPA with milder disease severity^14^. These findings highlight IPA as a potentially relevant component of the circulating metabolic response, although its increased abundance does not establish a causal contribution to the effects of butyrate on cardiac aging.

### Metabolomics profiling reveals extensive metabolic improvements upon butyrate supplementation

To extend these observations beyond the targeted organic acid panel, we performed broad plasma metabolomic profiling. The resulting profiles revealed heterogeneous responses across amino acid derivatives, aromatic compounds, nucleotide-related metabolites, sulfur-and one-carbon-associated compounds, tryptophan metabolites, and other biochemical classes (Figure 2D). Principal component analysis (PCA) showed differences in global metabolic profiles among young, aged control, and aged butyrate-treated mice, with the first two principal components accounting for 40.6% and 14.7% of total variance, respectively (Figure 2E). Classification of significantly altered metabolites by biochemical category identified widespread metabolic changes across these classes (Figure 2F). The numbers and directions of significant changes differed between aging-and treatment-associated comparisons, indicating that supplementation selectively remodeled multiple components of the circulating metabolome rather than uniformly reversing age-associated metabolic changes. Nucleotide-related metabolites exhibited a particularly consistent pattern. Guanosine, guanine, inosine, orotic acid, and cytidine were more abundant in aged control plasma than in young plasma and decreased following butyrate supplementation (Figure 2G). In contrast, hypoxanthine was less abundant in aged controls than in young mice and declined further following treatment. These findings suggest selective remodeling of circulating purine- and pyrimidine-associated metabolite pools.

Several metabolites associated with host–microbial co-metabolism and nitrogen handling also exhibited distinct age- and treatment-associated responses (Figure 2H). Indoxyl sulfate and hippurate were elevated in aged control plasma relative to young plasma and declined following butyrate supplementation. Conversely, tryptamine increased during aging and was further elevated after treatment, whereas valerate declined with aging and increased following supplementation. Urea and creatinine were also more abundant in aged controls than in young mice and decreased in treated aged mice. The circulating concentrations of these metabolites reflect contributions from microbial precursor production, host biotransformation, tissue metabolism, and excretion^10, 11^. Therefore, their changes cannot be attributed exclusively to altered gut microbial activity or interpreted as direct evidence of improved renal function but a beneficial response of many interacting systems. Additional plasma metabolite profiles revealed both treatment-associated shifts toward young-reference abundances and age-associated changes that persisted or became more pronounced following supplementation (Supplementary Figure 3). Together, these results reveal a heterogeneous but substantial reorganization of the circulating metabolome.

We next investigated whether butyrate supplementation was accompanied by changes in circulating lipid composition. Plasma lipidomic profiling revealed distinct abundance patterns across lysophosphatidylcholine (LPC), phosphatidylcholine (PC), phosphatidylethanolamine (PE), phosphatidylinositol (PI), phosphatidylserine (PS), hexosylceramide (HexCer), diacylglycerol (DAG), and triacylglycerol (TAG) species (Figure 2I). Several PC species exhibited higher abundance in aged controls than in young mice and lower abundance following butyrate supplementation. LPC (20:0) showed a similar age-associated increase and treatment-associated decrease. In contrast, selected plasmenyl-PE species were more abundant in treated aged mice, while individual PI and HexCer species also exhibited treatment-associated increases. TAG species showed more heterogeneous patterns, with several elevated during aging and displaying intermediate or reduced abundances after treatment. These findings indicate that late-life butyrate supplementation was associated with lipid-species-specific remodeling rather than uniform changes across lipid classes^19, 40,41^. Collectively, these analyses demonstrate that late-life sodium butyrate supplementation is associated with selective remodeling of the circulating metabolome and lipidome, encompassing central-carbon metabolism, hydroxybutyrates, nucleotide-related metabolites, host–microbial co-metabolites, and diverse lipid species.

### Late-life butyrate supplementation attenuates age-associated cardiac remodeling and functional decline

To determine whether the systemic metabolic improvements associated with late-life butyrate supplementation were accompanied by changes in cardiac aging, we assessed cardiac structure and function in young, aged control, and aged butyrate-treated C57BL/6J mice. Cardiac aging is characterized by progressive structural remodeling, myocardial hypertrophy, and impaired ventricular relaxation, which may develop alongside changes in systolic performance^3, 6, 7, 42, 43^. We therefore examined cardiac and pulmonary tissue mass together with echocardiographic indices of systolic and diastolic function.

LV mass normalized to tibial length was significantly higher in aged male controls than in young mice (P = 1.9 × 10^-5^; Figure 3A), consistent with age-associated cardiac hypertrophic remodeling. Butyrate-treated aged males exhibited significantly lower normalized LV mass than untreated aged controls (P = 4.7 × 10^-5^; Figure 3A), with values shifted toward the young-reference range. A similar pattern was observed in female mice, in which normalized LV mass increased with aging (P = 4.9 × 10^-5^) and was significantly lower following butyrate supplementation (P = 0.004; Supplementary Figure 4A). Lung mass normalized to tibial length was also higher in aged mice than in young mice in both sexes, whereas differences between treated and untreated aged groups were not statistically significant (Figure 3B and Supplementary Figure 4B). These findings indicate that butyrate supplementation attenuated age-associated LV hypertrophic remodeling without a corresponding significant reduction in normalized lung mass.

**Figure 3.**
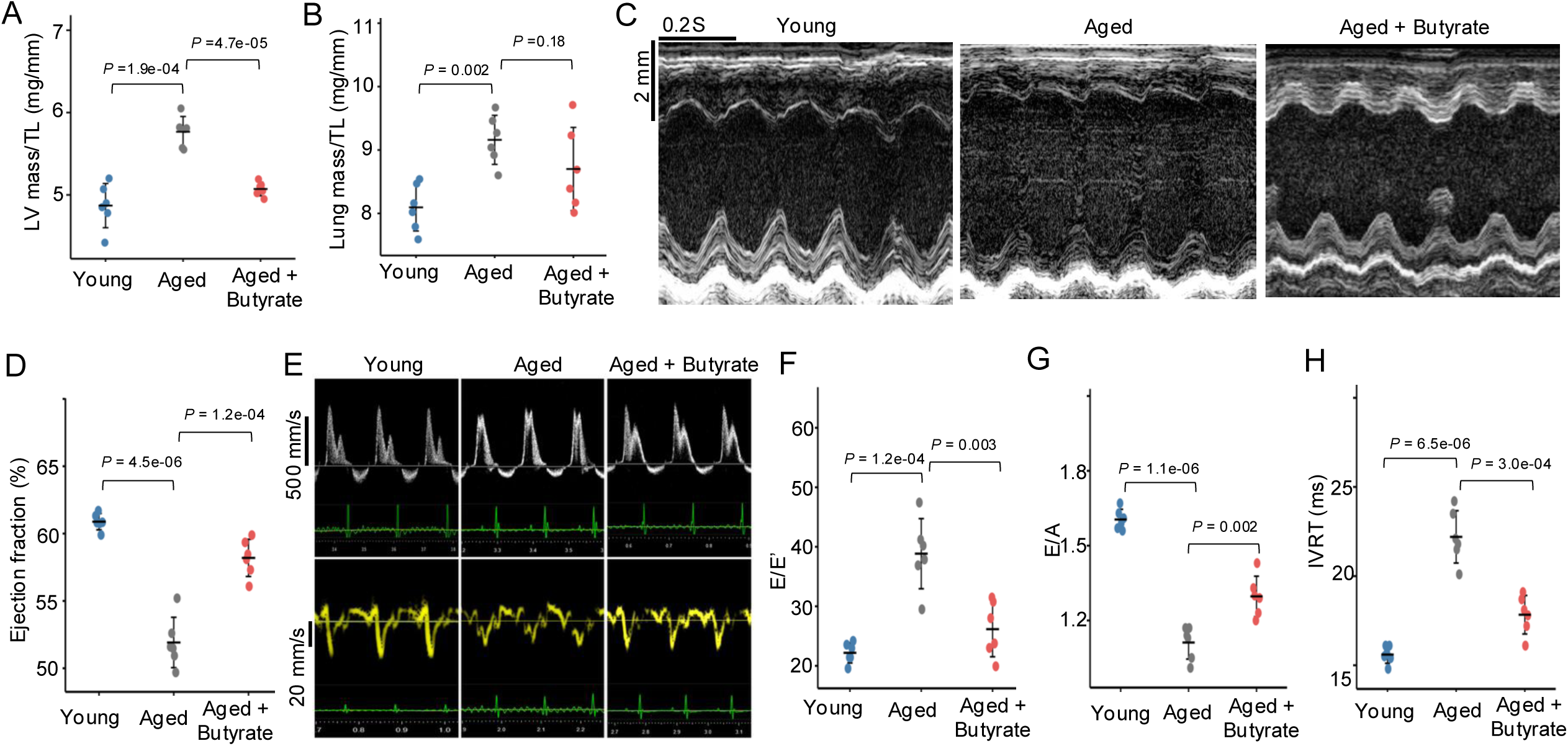
Late-life sodium butyrate supplementation attenuates age-associated cardiac hypertrophy and systolic and diastolic dysfunction in male mice. Cardiac structure and function were assessed in young (4-month-old), aged control (28-month-old), and aged sodium butyrate-treated (28-month-old) male C57BL/6J mice (n = 6 biologically independent mice per group). (A) LV mass normalized to TL, determined by weighing dissected LV tissue and expressed in mg/mm. (B) Lung mass normalized to tibial length (mg/mm). (C) Representative M-mode echocardiographic recordings illustrating LV wall motion and chamber dimensions in young, aged control, and aged butyrate-treated mice. Scale bars, 2 mm (vertical) and 0.2 s (horizontal). (D) LV ejection fraction (%) determined by transthoracic echocardiography. (E) Representative transmitral pulsed-wave Doppler recordings (upper panels) and mitral annular tissue Doppler recordings (lower panels) in young, aged control, and aged butyrate-treated mice. Velocity scale bars, 500 mm/s for transmitral Doppler and 20 mm/s for tissue Doppler. (F) Ratio of E/E′. (G) Ratio of E/A. (H) IVRT, measured as the interval between aortic valve closure and mitral valve opening and expressed in milliseconds. Each point represents an individual mouse; horizontal summary marks and error bars indicate mean ± SD. Between-group comparisons were performed using two-sided, independent, equal-variance Student’s t-tests with BH correction. Young reference animals were 4 months old, and aged animals were assessed at 28 months. Brackets indicate comparisons between young and aged control mice and between aged control and aged butyrate-treated mice. Displayed P values were adjusted using the Benjamini–Hochberg procedure within the prespecified multiple-comparison families. E, early transmitral filling velocity; A, late transmitral filling velocity; E′, early diastolic mitral annular velocity; LV, left ventricle; TL, tibial length; IVRT, isovolumic relaxation time. Doppler-derived indices were interpreted collectively as measures of diastolic function rather than as direct measurements of LV filling pressure.

We next evaluated LV function using echocardiography. Representative M-mode recordings illustrated differences in ventricular wall motion and chamber dimensions among young, aged control, and aged butyrate-treated male mice (Figure 3C). Quantitative analysis showed that LV ejection fraction was significantly lower in aged controls than in young mice (P = 4.5 × 10^-6^; Figure 3D), whereas treated aged males exhibited significantly higher ejection fraction than untreated aged controls (P = 1.2 × 10^-4^). Although systolic function is often relatively preserved during earlier stages of cardiac aging, previous studies have reported reductions in EF at advanced ages, including a significant decline in C57BL/6J mice at 29–31 months of age^42^ and a modest reduction in other mouse cohorts at approximately 30 months^44^, consistent with our findings in 28-month-old mice. Female mice exhibited a similar pattern, with ejection fraction decreasing during aging (P = 4.5 × 10^-4^) and increasing following butyrate supplementation (P = 0.008; Supplementary Figure 4C). These results suggest that late-life butyrate supplementation attenuated the age-associated decline in LV systolic performance in both sexes.

Because impaired ventricular relaxation is a prominent feature of cardiac aging, we further assessed diastolic function using pulsed wave transmitral Doppler and tissue Doppler imaging^7, 8, 27^. Representative recordings demonstrated age- and treatment-associated differences in mitral inflow and annular motion profiles (Figure 3E). The ratio of early transmitral filling velocity to early diastolic mitral annular velocity (E/E′) was significantly higher in aged male controls than in young mice (P = 1.2 × 10^-4^), whereas butyrate-treated aged males exhibited a significantly lower E/E′ ratio than untreated aged controls (P = 0.003; Figure 3F). Female mice exhibited a similar pattern, with E/E′ increasing during aging (P = 1.4 × 10^-4^) and decreasing following butyrate supplementation (P = 8.3 × 10^-4^; Supplementary Figure 4D).

The ratio of early to late transmitral filling velocities (E/A) was significantly lower in aged male controls than in young mice (P = 1.1 × 10^-6^; Figure 3G), indicating an age-associated alteration in the transmitral filling pattern. Butyrate supplementation significantly increased E/A relative to aged controls (P = 0.002). Female mice showed corresponding differences, with E/A decreasing during aging (P = 3.1 × 10^-6^) and increasing following treatment (P = 0.002; Supplementary Figure 4E). The concordant treatment-associated changes in E/E′ and E/A suggest a shift in the diastolic filling profile toward that observed in young mice.

To further assess LV relaxation, we measured isovolumic relaxation time (IVRT), the interval between aortic valve closure and mitral valve opening. IVRT was significantly prolonged in aged male controls compared with young mice (P = 6.5 × 10^-6^), whereas butyrate supplementation was associated with a significantly shorter IVRT relative to untreated aged controls (P = 3.0 × 10^-4^; Figure 3H). Female mice exhibited the same directional pattern, with IVRT increasing during aging (P = 4.9 × 10^-6^) and decreasing following treatment (P = 1.2 × 10^-4^; Supplementary Figure 4F). These findings provide additional evidence that butyrate supplementation attenuated age-associated abnormalities in ventricular relaxation. Collectively, the reductions in normalized LV mass and the favorable changes in ejection fraction, transmitral filling indices, and IVRT indicate that late-life butyrate supplementation attenuates multiple features of age-associated cardiac structural and functional decline in both male and female mice.

### Late-life butyrate supplementation remodels myocardial metabolism, extracellular matrix, and protein homeostasis

To investigate the molecular changes accompanying the attenuation of age-associated cardiac remodeling and functional decline, we performed integrated metabolomic, lipidomic, and proteomic analyses of LV tissue from young, aged control, and aged butyrate-treated male mice. Myocardial metabolomic profiling revealed distinct age- and treatment-associated changes across metabolites involved in amino acid metabolism, intermediary metabolism, and host–microbial co-metabolism (Figure 4A and B). Several metabolites, including γ-aminobutyric acid (GABA), N-acetylproline, and homocysteine thiolactone, were more abundant in aged control hearts than in young hearts and decreased following butyrate supplementation. In contrast, pyroglutamic acid and pipecolic acid were less abundant in aged controls and remained comparatively low in treated aged mice. Other metabolites displayed responses that further accentuated their age-associated differences. Ornithine, kynurenine, and methionine sulfoxide increased with aging and were further elevated following treatment, whereas hypotaurine and methylmalonic acid decreased with aging and declined further in hearts from treated animals (Figure 4B). PCA and additional myocardial metabolite profiles supported substantial age-associated metabolic differences, with heterogeneous responses to supplementation (Supplementary Figure 5A and B). Together, these findings indicate that late-life butyrate treatment selectively remodels myocardial metabolite pools rather than uniformly restoring a young metabolic profile, consistent with the tissue-dependent nature of metabolic aging^20^.

**Figure 4.**
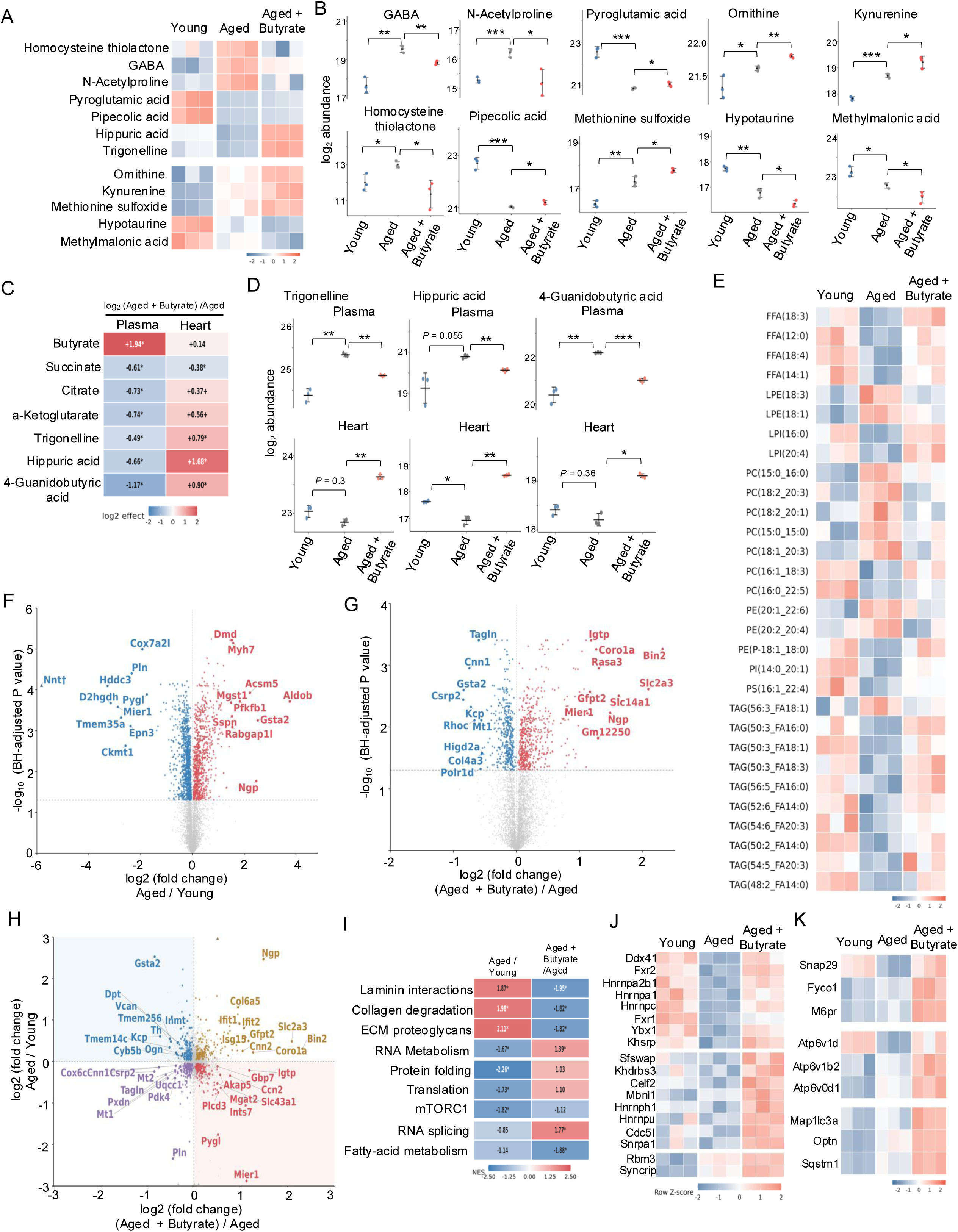
Late-life sodium butyrate supplementation is associated with compartment-dependent myocardial metabolic remodeling and selective changes in extracellular-matrix, RNA-processing, and autophagy-associated protein programs. LV metabolomic, lipidomic, and proteomic analyses were performed in young (4-month-old), aged control (28-month-old), and aged sodium butyrate-treated (28-month-old) male C57BL/6J mice (n = 3 biologically independent mice per group for each omics dataset). (A) Heatmap of selected LV metabolites associated with intermediary metabolism, amino acid metabolism, and host–microbial co-metabolism. Rows indicate metabolites and columns represent individual mice. Colors indicate row-standardized relative abundance. (B) Representative LV metabolite abundances, including γ-aminobutyric acid (GABA), N-acetylproline, pyroglutamic acid, ornithine, kynurenine, homocysteine thiolactone, pipecolic acid, methionine sulfoxide, hypotaurine, and methylmalonic acid. Abundances are displayed on a log₂ scale. Brackets indicate young versus aged control and aged control versus aged butyrate-treated comparisons. (C) Heatmap comparing treatment-associated changes in selected metabolites measured in plasma and LV tissue. Values indicate log₂ fold changes for aged butyrate-treated versus aged control mice within each compartment. Positive and negative values indicate higher and lower abundance following treatment, respectively. Asterisks denote statistically significant changes after multiple-testing correction. (D) Plasma and LV abundance profiles of trigonelline, hippuric acid, and 4-guanidinobutyric acid in young, aged control, and aged butyrate-treated mice. Upper panels show plasma measurements, and lower panels show myocardial measurements. Lines connect group means across independent experimental groups and do not represent longitudinal measurements of individual animals. Abundances are displayed on a log₂ scale. (E) Heatmap of selected LV lipid species, including free fatty acids (FFAs), lysophosphatidylethanolamines (LPEs), lysophosphatidylinositols (LPIs), phosphatidylcholines (PCs), phosphatidylethanolamines (PEs), phosphatidylinositols (PIs), phosphatidylserines (PSs), and triacylglycerols (TAGs). Colors indicate row-standardized relative abundance within each lipid species. (F and G) Volcano plots showing differential left ventricular protein abundance in aged control versus young mice (F) and aged butyrate-treated versus aged control mice (G). The horizontal axis represents log₂ fold change, and the vertical axis represents −log₁₀(Benjamini–Hochberg-adjusted P value). Red and blue points indicate proteins with significantly higher and lower abundance, respectively, in the numerator group, defined by an adjusted false-discovery rate (FDR) < 0.05. Selected differentially abundant proteins are labeled. (H) Comparison of age-associated and treatment-associated LV protein log₂ fold changes. The horizontal axis represents aged butyrate-treated versus aged control mice, and the vertical axis represents aged control versus young mice. Proteins in the upper-left and lower-right quadrants exhibit treatment-associated changes opposing the direction of their age-associated changes. The other quadrants indicate concordant directions of change. (I) Normalized enrichment scores (NESs) for selected pathways identified using Reactome, Gene Ontology Biological Process, and Hallmark gene-set collections. Comparisons include aged control versus young and aged butyrate-treated versus aged control mice. Positive and negative NES values indicate enrichment in opposite directions. Asterisks indicate pathways meeting the specified FDR threshold after correction across the tested gene-set collection. (J) Heatmap of selected LV proteins associated with RNA metabolism, RNA binding, and pre-mRNA processing, including heterogeneous nuclear ribonucleoproteins and other RNA-processing factors. (K) Heatmap of selected LV proteins associated with autophagy, lysosomal function, and intracellular vesicle trafficking, including SNAP29, FYCO1, M6PR, ATP6V1D, ATP6V1B2, ATP6V0D1, MAP1LC3A, OPTN, and SQSTM1. Individual points in metabolite abundance plots represent biologically independent mice; summary marks and error bars indicate mean ± SD. Heatmap columns represent independent biological samples, and row-standardized colors indicate relative abundance within each molecular feature. For individual metabolite comparisons, displayed P values were adjusted using the Benjamini–Hochberg procedure within the prespecified comparison families. Protein differential-abundance significance was defined as BH-adjusted FDR < 0.05. Pathway significance was assessed using FDR correction across the corresponding tested gene-set collections. * *P* < 0.05; ** *P* < 0.01; *** *P* <0.001.

We next compared metabolites measured in both plasma and LV tissue to determine whether circulating responses reflected corresponding myocardial changes (Figure 4C). Despite the substantial increase in circulating butyrate, its myocardial abundance changed only modestly. Succinate decreased in both plasma and myocardium following supplementation, whereas citrate and α-ketoglutarate exhibited opposing responses, decreasing in plasma but increasing in LV tissue. These observations demonstrate that changes in circulating metabolite abundance do not necessarily predict corresponding myocardial responses, highlighting compartment-dependent metabolic remodeling.

Three metabolites, including trigonelline, hippuric acid, and 4-guanidinobutyric acid (4-GBA), displayed particularly striking reciprocal changes between plasma and myocardium (Figure 4C and D). All three were more abundant in plasma from aged control mice than from young mice and decreased following butyrate supplementation. In contrast, their myocardial abundances were lower in aged controls than in young hearts and increased following treatment. These findings further highlight the compartment-dependent nature of metabolic remodeling during aging and supplementation. Trigonelline is of particular interest because it has been linked to nicotinamide adenine dinucleotide (NAD⁺) metabolism and muscle function during aging^45^, with additional studies investigating its metabolic effects in HFpEF^46^. Hippuric acid is a host– microbial co-metabolite whose circulating abundance reflects microbial precursor production, host conjugation, and clearance^10, 11^.

LV lipidomic profiling further revealed selective remodeling of myocardial lipid species (Figure 4E). Several free fatty acids, including FFA (18:3), FFA (12:0), FFA (18:4), and FFA (14:1), exhibited lower abundance in aged control hearts than in young hearts and increased following butyrate supplementation. Selected lysophospholipids, including LPI (16:0) and LPI (20:4), showed similar directional responses. In contrast, phosphatidylcholine (PC) and phosphatidylethanolamine (PE) species displayed heterogeneous patterns, with individual molecular species exhibiting distinct age-and treatment-associated changes. Several triacylglycerol (TAG) species were less abundant in aged hearts than in young hearts, with variable responses to treatment. These findings indicate that butyrate supplementation selectively alters myocardial lipid composition rather than uniformly affecting entire lipid classes.

### Late-life butyrate supplementation attenuates myocardial fibrosis and remodels extracellular-matrix and protein homeostasis pathways

To identify protein programs associated with myocardial metabolic and functional remodeling, we performed proteomic profiling of LV tissue. Aging was associated with extensive changes in protein abundance, with 1,843 differentially abundant proteins between aged control and young hearts at a Benjamini–Hochberg-adjusted false-discovery rate (FDR) below 0.05, including 833 proteins with higher and 1,010 with lower abundance in aged hearts (Figure 4F and Supplementary Figure 6A). Comparison of butyrate-treated and untreated aged hearts identified 1,008 differentially abundant proteins, including 554 with higher and 454 with lower abundance following treatment (Figure 4G and Supplementary Figure 6A). An additional comparison between treated aged and young hearts identified 2,204 differentially abundant proteins, indicating that the treated myocardial proteome remained distinct from the young-reference state. Comparison of age-associated and treatment-associated protein fold changes revealed subsets of proteins whose responses to supplementation opposed their age-associated direction (Figure 4H), consistent with selective remodeling rather than global reversal of proteomic aging.

Pathway enrichment analysis identified ECM remodeling among the prominent treatment-associated responses (Figure 4I and Supplementary Figure 6B and C). Reactome pathways associated with laminin interactions, collagen degradation, and ECM proteoglycans were positively enriched in aged hearts relative to young controls but negatively enriched in butyrate-treated aged hearts relative to untreated aged controls. These opposing enrichment patterns indicate that supplementation altered selected age-associated ECM-related protein programs. To examine whether these molecular changes were accompanied by histological differences, we performed picrosirius red staining of aged control and treated hearts. Representative sections revealed differences in perivascular and interstitial collagen deposition (Supplementary Figure 7A). Quantification demonstrated a significantly lower collagen-positive myocardial area in butyrate-treated aged hearts than in untreated aged controls (P = 0.002; Supplementary Figure 7B). These findings provide complementary histological evidence of attenuated myocardial fibrotic remodeling.

In addition to ECM remodeling, proteomic analysis identified prominent changes in RNA-processing pathways. Metabolism of RNA pathway was negatively enriched in aged hearts relative to young hearts and positively enriched following butyrate supplementation, with significant enrichment in both comparisons (Figure 4I and Supplementary Figure 6C). RNA-splicing-associated proteins also contributed to the treatment-associated response. Representative protein profiles revealed changes in multiple RNA-binding and RNA-processing factors, including heterogeneous nuclear ribonucleoproteins and other components of the splicing machinery (Figure 4J). These observations identify RNA metabolism and processing as prominent components of myocardial proteomic remodeling and motivated the subsequent full-length transcriptomic analysis.

Pathways associated with protein folding, translation, and mTORC1 signaling displayed distinct responses. Because mTORC1 signaling is functionally connected to autophagy regulation, which is important for cardiac homeostasis^5, 21–23^, we further examined proteins involved in autophagosome formation, vesicular trafficking, and lysosomal function. Proteomic profiles revealed changes in selected proteins, including SNAP29, FYCO1, M6PR, ATP6V1D, ATP6V1B2, ATP6V0D1, MAP1LC3A, OPTN, and SQSTM1 (Figure 4K and Supplementary Figure 6D–G). Several exhibited lower abundances during aging and higher abundance following supplementation, whereas others displayed distinct response patterns. These findings indicate heterogeneous remodeling of autophagy-associated protein machinery rather than a uniform response across the pathway.

Collectively, these analyses reveal extensive but selective myocardial molecular remodeling following late-life sodium butyrate supplementation. Compartment-dependent metabolite responses distinguish myocardial changes from those observed in plasma, while lipidomics identifies species-specific alterations in cardiac lipid composition. Proteomic and histological analyses further demonstrate attenuation of ECM-associated remodeling and myocardial fibrosis, together with changes in RNA-processing and autophagy-associated protein programs. These findings provide a molecular framework for the subsequent investigation of transcript-specific regulation and mitochondrial quality-control pathways.

### Late-life butyrate supplementation remodels cardiac transcript expression and autophagy-associated RNA programs

The proteomic identification of RNA metabolism and splicing-associated pathways prompted us to investigate whether late-life butyrate supplementation also altered myocardial transcript expression and isoform composition. To resolve these changes beyond conventional gene-level measurements, we performed PacBio Kinnex full-length RNA sequencing of LV tissue from young, aged control, and aged butyrate-treated male mice (Figure 5A). Long-read transcript sequencing enables the characterization of full-length transcript structures and the quantification of individual RNA isoforms, providing additional resolution for studying transcriptomic remodeling during cardiac aging^24, 28, 29^.

**Figure 5.**
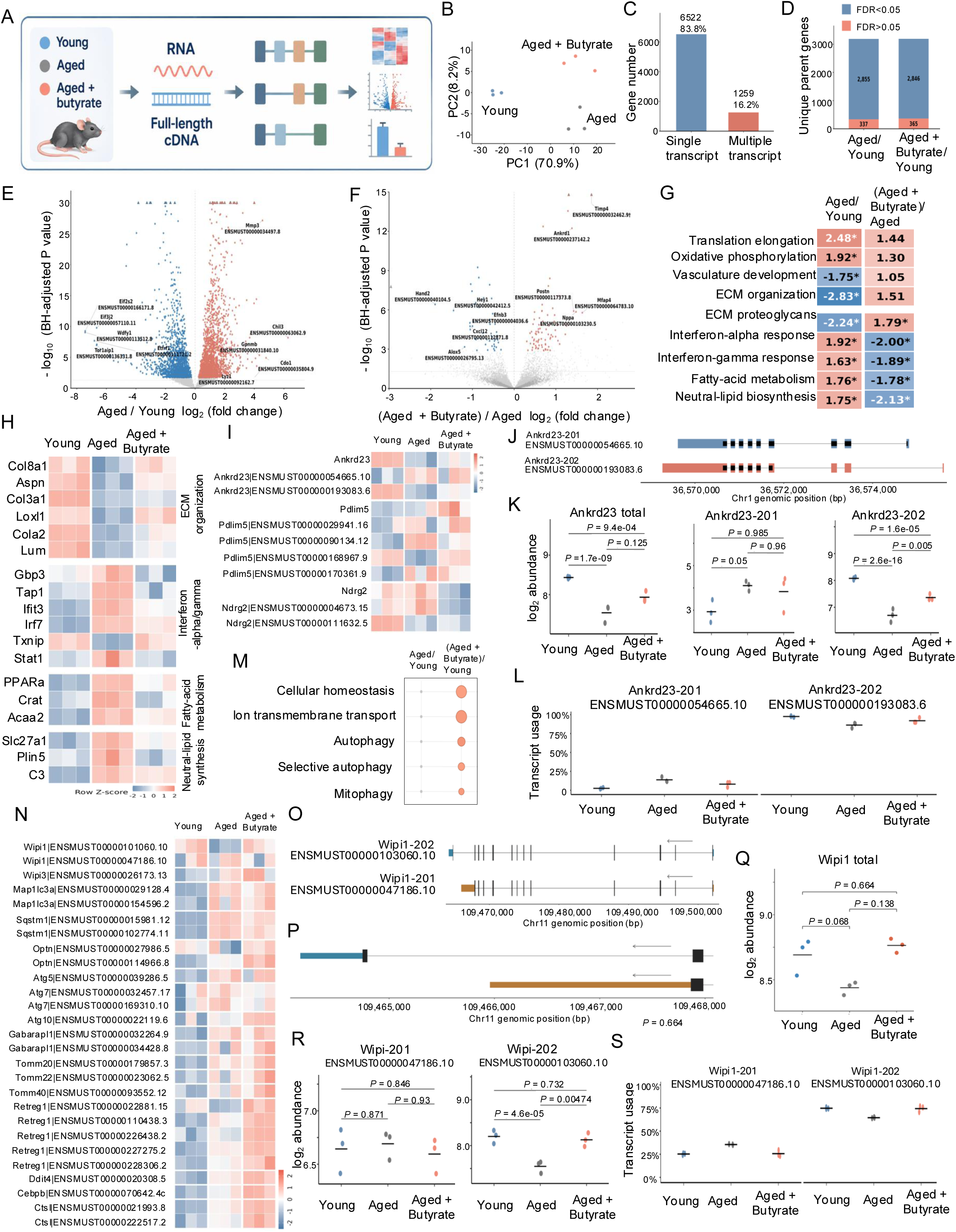
Full-length RNA sequencing reveals selective transcriptomic remodeling and autophagy-associated transcript changes following late-life sodium butyrate supplementation. PacBio Kinnex full-length RNA sequencing was performed on left ventricular (LV) tissue from young (4-month-old), aged control (28-month-old), and aged sodium butyrate-treated (28-month-old) male C57BL/6J mice (n = 3 biologically independent mice per group). (A) Schematic illustrating the experimental workflow for full-length RNA sequencing, including RNA isolation from LV tissue, full-length cDNA preparation, transcript characterization, and downstream gene-and transcript-level analyses. (B) Principal component analysis (PCA) of LV RNA-expression profiles across the three experimental groups. Each point represents an independent mouse. Principal components 1 and 2 explain 70.9% and 8.2% of total variance, respectively. (C) Numbers of genes represented by a single detected transcript (6,522) or multiple detected transcripts (1,259) in the full-length RNA-sequencing dataset. (D) Numbers of parent genes containing at least one significantly differentially expressed transcript despite not reaching the corresponding gene-level significance threshold. In the aged-versus-young comparison, 337 parent genes met this criterion; in the aged-butyrate-versus-young comparison, 365 parent genes met this criterion. Blue and salmon segments indicate parent genes with FDR < 0.05 and FDR > 0.05, respectively, according to the figure annotation. (E and F) Volcano plots showing differential transcript abundance in aged control versus young (E) and aged butyrate-treated versus aged control (F) mice. The horizontal axis represents transcript-level log₂ fold change, and the vertical axis represents −log₁₀ P value. Blue and salmon points indicate transcripts with lower and higher abundance, respectively, in the numerator group. Selected transcripts and their corresponding genes are annotated. (G) Normalized enrichment scores (NESs) for selected biological pathways in aged control versus young and aged butyrate-treated versus aged control comparisons. Pathways include translation elongation, oxidative phosphorylation, vasculature development, extracellular-matrix organization, ECM proteoglycans, interferon-α response, interferon-γ response, fatty-acid metabolism, and neutral-lipid biosynthesis. Positive and negative NES values indicate enrichment in opposite directions. Asterisks denote pathways meeting the specified multiple-testing-adjusted significance threshold. (H) Row-standardized heatmaps of selected transcripts associated with ECM organization, interferon-α/γ response, and fatty-acid metabolism. Each column represents an individual mouse, and colors indicate relative transcript abundance within each feature. (I) Heatmap of parent-gene and transcript-level expression profiles for Ankrd23, Pdlim5, and Ndrg2. Individual annotated transcript identifiers are displayed alongside their corresponding parent genes. (J) Genomic structures of annotated Ankrd23-201 (ENSMUST00000054665.10) and Ankrd23-202 (ENSMUST00000193083.6) transcripts, illustrating differences in exon organization. (K) Total Ankrd23 gene abundance and individual Ankrd23-201 and Ankrd23-202 transcript abundances across the three experimental groups. Abundances are displayed on a log₂ scale. (L) Relative transcript representation of Ankrd23-201 and Ankrd23-202, expressed as the percentage of the combined detected Ankrd23 transcript abundance. (M) Functional-category analysis highlighting cellular homeostasis, ion transmembrane transport, autophagy, selective autophagy, and mitophagy among the represented transcript-associated biological processes. Bubble size indicates the corresponding category representation as defined by the enrichment analysis. (N) Row-standardized heatmap of selected transcripts associated with autophagy, lysosomal function, membrane trafficking, and mitochondrial quality control, including transcripts from Wipi1, Wipi3, Map1lc3a, Sqstm1, Optn, Atg5, Atg7, Atg10, Gabarapl1, Tomm20, Tomm22, Tomm40, Retreg1, Ddit4, Cebpb, and Ctsl. (O and P) Annotated transcript structures of Wipi1-202 (ENSMUST00000103060.10) and Wipi1-201 (ENSMUST00000047186.10), shown at different genomic scales to illustrate transcript-specific exon organization. (Q) Total Wipi1 gene abundance across young, aged control, and aged butyrate-treated mice. (R) Abundances of individual Wipi1-201 and Wipi1-202 transcripts across the three groups, displayed on a log₂ scale. (S) Relative transcript representation of Wipi1-201 and Wipi1-202, expressed as the percentage of the combined detected Wipi1 transcript abundance. Gene-and transcript-level differential-expression analyses were performed using DESeq2, with BH-adjusted P values and independent filtering. Significance required an available adjusted P < 0.05 and a finite effect estimate. PCA was performed on variance-stabilized expression values. Transcript-level differential expression was not interpreted as direct evidence of differential transcript usage.

PCA revealed distinct cardiac transcript-expression profiles among the three groups, with the first two principal components explaining 70.9% and 8.2% of the variance, respectively (Figure 5B). Young and aged control samples occupied different regions of the expression space, whereas butyrate-treated aged samples formed a separate group, indicating substantial age-and treatment-associated transcriptomic differences. Full-length RNA sequencing retained 9,353 transcripts corresponding to 7,781 parent genes, including 6,522 genes represented by a single retained transcript (83.8%) and 1,259 genes represented by multiple retained transcripts (16.2%; Figure 5C). In parallel, gene-level differential-expression analysis included 9,421 genes, of which 1,640 lacked corresponding retained transcripts in the transcript-level dataset. Differential-expression analysis revealed age-and treatment-associated transcript-level changes (Figure 5D–F). Notably, 337 parent genes in the aged-versus-young comparison and 365 parent genes in the treated-aged-versus-aged comparison contained at least one significantly differentially expressed transcript despite not reaching the corresponding gene-level significance threshold (Figure 5D). These findings demonstrate that transcript-level analysis captures expression differences that may not be apparent from aggregated parent-gene measurements.

Gene-set enrichment analysis identified selective changes in biological programs associated with aging and butyrate supplementation (Figure 5G). Interferon-α and interferon-γ response signatures were positively enriched during aging and negatively enriched following treatment. These findings indicate that treatment opposed selected age-associated interferon-response signatures, consistent with previous observations linking inflammatory and interferon-associated transcriptional programs to myocardial aging^25, 26^. Fatty-acid metabolism and neutral-lipid biosynthesis exhibited similar opposing enrichment patterns, suggesting treatment-associated changes in myocardial lipid-related transcriptional programs that complement the lipidomic findings in Figure 4. In contrast, ECM-associated pathways showed heterogeneous responses: ECM proteoglycan pathways were negatively enriched with aging and positively enriched following treatment, whereas broader ECM organization and vasculature-development pathways exhibited age-associated changes without significant treatment-associated enrichment. Translation elongation and oxidative phosphorylation also showed significant age-associated enrichment without corresponding significant opposing responses following supplementation. Together, these results indicate that butyrate selectively remodeled inflammatory, metabolic, and ECM-associated transcriptional programs rather than uniformly reversing age-associated expression changes.

Representative expression profiles further illustrated these pathway-specific responses (Figure 5H). ECM-associated genes, including Col8a1, Aspn, Col3a1, Loxl1, Col1a2, and Lum, showed lower transcript abundance in aged hearts and higher abundance following treatment. Conversely, multiple interferon-responsive genes, including Gbp3, Tap1, Ifit3, Irf7, and Txnip, exhibited higher expression in aged hearts and lower expression in treated aged hearts. Selected fatty-acid-metabolism genes, including PPARa, Crat, Acaa2, Slc27a1, and Plin5, also displayed treatment-associated changes. These heterogeneous RNA-level patterns complement the protein-level and lipidomic results described above.

We next examined individual loci to determine how transcript-level measurements differed from aggregated parent-gene expression. Three representative genes, Ankrd23, Pdlim5, and Ndrg2, displayed distinct transcript-specific abundance patterns across the experimental groups (Figure 5I). At the Ankrd23 locus, full-length sequencing distinguished two annotated transcripts, Ankrd23-201 and Ankrd23-202, with distinct transcript structures (Figure 5J). Total Ankrd23 expression decreased significantly during aging and increased following supplementation (Figure 5K). The predominant transcript, Ankrd23-202, broadly followed the parent-gene expression pattern, whereas Ankrd23-201 displayed a different distribution across groups (Figure 5K and L). These findings illustrate the additional resolution provided by transcript-level analysis without necessarily implying altered relative isoform usage.

Functional-category analysis further highlighted cellular homeostasis, ion transmembrane transport, autophagy, selective autophagy, and mitophagy among treatment-associated transcriptional changes (Figure 5M), consistent with the proteomic remodeling of autophagy-related proteins in Figure 4. We therefore examined genes involved in autophagosome formation, lysosomal trafficking, and mitochondrial quality control. Transcript-level profiles revealed heterogeneous expression patterns involving Wipi1, Wipi3, Map1lc3a, Sqstm1, Optn, Atg5, Atg7, Atg10, Gabarapl1, Tomm20, Tomm22, Tomm40, Retreg1, Ddit4, Cebpb, and Ctsl (Figure 5N). Several transcripts showed lower abundance during aging and higher abundance following supplementation, whereas others followed distinct patterns.

Among these candidates, Wipi1 exhibited a particularly notable transcript-specific response. Full-length sequencing distinguished two annotated transcripts, Wipi1-201 and Wipi1-202, with distinct transcript structures (Figure 5O and P). Although total Wipi1 abundance did not differ significantly among young, aged control, and aged butyrate-treated hearts (Figure 5Q), Wipi1-202 abundance decreased significantly with aging and increased significantly following supplementation (Figure 5R). In contrast, Wipi1-201 showed no significant changes in the corresponding comparisons. Transcript representation analysis indicated that Wipi1-202 was the predominant detected isoform across groups, accounting for approximately two-thirds to three-quarters of total Wipi1 transcript abundance (Figure 5S). These findings identify an age-and treatment-responsive Wipi1 transcript whose differential abundance was not captured by parent-gene-level significance testing.

WIPI1 is a phosphatidylinositol-3-phosphate-binding protein associated with autophagic membrane formation and early autophagosome biogenesis^47, 48^. WIPI-family proteins participate in the recruitment and organization of molecular machinery required for autophagosome formation, functionally connecting phosphoinositide signaling to the LC3/GABARAP conjugation systems involving ATG7 and related autophagy proteins^5, 23, 48, 49^. Previous studies have also implicated WIPI1 in cardiac disease and mitochondrial quality-control processes, including experimental ventricular dysfunction and diabetic cardiomyopathy^50, 51^. The treatment-associated increase in Wipi1-202 therefore identifies a candidate transcript potentially relevant to the cardiac autophagy response. Collectively, these results demonstrate that late-life butyrate supplementation induces extensive but selective myocardial transcriptional remodeling, including changes in inflammatory, metabolic, ECM-associated, and autophagy-related RNA programs.

### Late-life butyrate supplementation preserves cardiac mitochondrial ultrastructure and respiratory function

Mitochondrial dysfunction and impaired organelle quality control are prominent features of cardiac aging and contribute to declining myocardial bioenergetic capacity^3, 52–56^. Given the treatment-associated changes in myocardial metabolism and autophagy-related molecular programs identified by multi-omics analyses, we next investigated whether late-life butyrate supplementation was accompanied by preservation of mitochondrial structure and respiratory function in the aging heart. We examined LV mitochondrial ultrastructure using transmission electron microscopy (TEM) and measured oxygen consumption in isolated cardiac mitochondria from young, aged control, and aged butyrate-treated mice (Figure 6).

**Figure 6.**
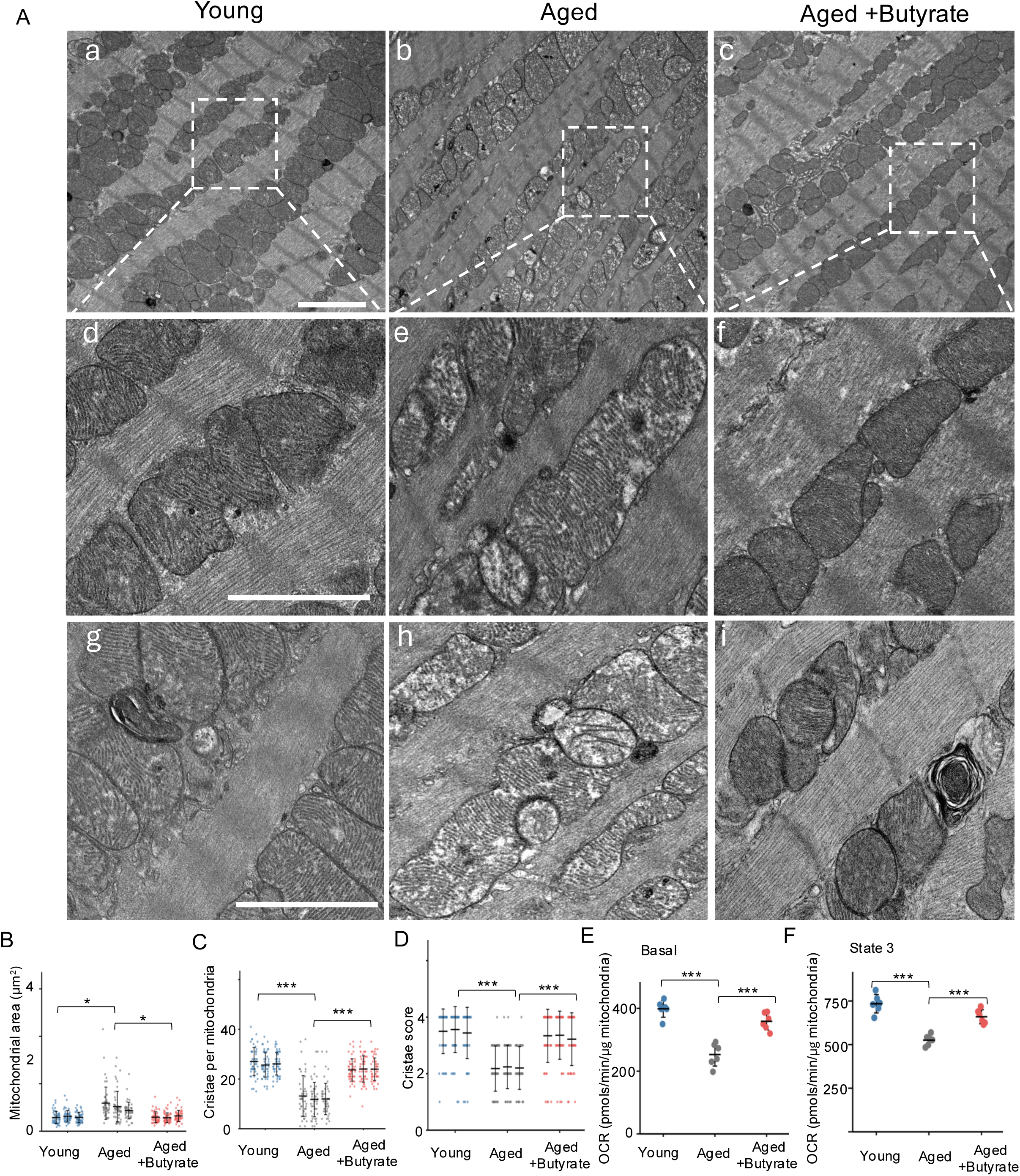
Late-life sodium butyrate supplementation attenuates age-associated mitochondrial ultrastructural abnormalities and improves cardiac mitochondrial respiration. Mitochondrial ultrastructure and respiratory function were assessed in young (4-month-old), aged control (28-month-old), and aged sodium butyrate-treated (28-month-old) male C57BL/6J mice. (A) Representative transmission electron microscopy (TEM) images of left ventricular myocardium from young, aged control, and aged butyrate-treated mice. Images a–c show lower-magnification views of cardiomyocyte mitochondrial organization, with dashed boxes indicating regions enlarged in images d–f, respectively. Images g–i show additional higher-magnification examples of mitochondrial ultrastructure and membrane-associated structures, with arrows indicating selected membranous profiles. Scale bar = 2µm. Aged control hearts exhibit enlarged mitochondrial profiles and disrupted cristae organization, whereas treated aged hearts display comparatively preserved mitochondrial morphology and cristae architecture. (B) Mitochondrial cross-sectional area (μm²) quantified from two-dimensional TEM sections. Measurements represent mitochondrial profile area rather than mitochondrial volume. (C) Number of discernible cristae per mitochondrial profile, quantified from TEM images. (D) Semiquantitative mitochondrial cristae morphology score, with higher scores indicating more intact and organized cristae architecture according to the scoring criteria described in STAR Methods. For panels B–D, three biologically independent mice were analyzed per group, with 50 mitochondrial profiles evaluated per mouse for each endpoint. Individual mitochondrial measurements are displayed within mouse-specific clusters, with horizontal summary lines and error bars indicating mean ± SD for each mouse. Statistical comparisons were performed using mouse-level summary measurements rather than treating individual mitochondrial profiles as independent biological replicates. (E) Substrate-supported baseline oxygen consumption rate (OCR) in isolated cardiac mitochondria measured using a Seahorse extracellular flux analyzer. Baseline OCR was recorded in the presence of pyruvate and malate before ADP injection. (F) ADP-stimulated State 3 OCR in isolated cardiac mitochondria, measured following ADP administration in the presence of pyruvate and malate. For panels E and F, OCR was normalized to mitochondrial protein and expressed as pmol O₂/min/μg mitochondrial protein. Each point represents one biologically independent mouse (n = 6 per group), with technical replicate wells averaged to obtain one value per mouse. Horizontal summary lines and error bars indicate mean ± SD. Young mice are shown in blue, aged controls in gray, and butyrate-treated aged mice in salmon. Between-group comparisons were performed using two-sided, independent, equal-variance Student’s t-tests with Benjamini–Hochberg correction. Brackets indicate the adjusted P values for aged control versus young and aged butyrate-treated versus aged control. * *P* < 0.05; ** *P* < 0.01; *** *P* <0.001.

TEM revealed pronounced age-associated alterations in mitochondrial morphology and internal membrane organization (Figure 6A). Young cardiomyocytes exhibited mitochondria with relatively compact morphology and well-organized cristae. In contrast, aged control hearts displayed accumulation of enlarged and damaged mitochondria with cristae disruption. These ultrastructural abnormalities are consistent with previous studies documenting mitochondrial enlargement, altered cristae architecture, and deterioration of mitochondrial organization during cardiac aging^52–56^. Notably, mitochondria in butyrate-treated aged hearts exhibited more compact structure and preserved cristae organization than those in untreated aged hearts, with ultrastructural features more closely resembling those observed in young controls (Figure 6A).

We next quantified mitochondrial area in TEM sections. Aged control hearts exhibited significantly larger mitochondria than young hearts (P = 0.013; Figure 6B), consistent with age-associated mitochondrial enlargement. In contrast, butyrate-treated aged hearts showed significantly smaller mitochondrial cross-sectional areas than aged controls (P = 0.013). We next quantified mitochondrial cristae organization, a structural determinant of inner mitochondrial membrane architecture and oxidative phosphorylation^53, 54^. Cristae number per mitochondria was markedly reduced in aged control hearts compared with young hearts (Figure 6C). However, butyrate supplementation significantly increased cristae number relative to untreated aged controls (P = 4.7 × 10^-5^; Figure 6C), indicating preservation of internal mitochondrial membrane organization. Consistent with these findings, assessment of cristae morphology revealed significantly lower scores in aged hearts than in young controls (P = 4.4 × 10^-5^), whereas butyrate-treated aged hearts exhibited higher scores than untreated aged hearts (P = 4.7 × 10^-5^; Figure 6D). These morphometric measurements demonstrate that late-life butyrate supplementation was associated with attenuation of age-related mitochondrial enlargement and cristae disorganization.

To determine whether the structural differences were accompanied by changes in mitochondrial respiratory capacity, we measured oxygen consumption rate (OCR) in isolated cardiac mitochondria using a Seahorse extracellular flux analyzer. Respiration was assessed using pyruvate and malate as substrates, with measurements obtained before and after ADP administration. Baseline OCR was significantly lower in mitochondria isolated from aged control hearts than in those from young hearts (P = 4.1 × 10^-5^; Figure 6E). However, butyrate supplementation significantly increased baseline OCR relative to aged controls (P = 3.0 × 10^-4^). We next assessed ADP-stimulated State 3 respiration, which reflects oxygen consumption during ADP-supported oxidative phosphorylation. State 3 OCR was significantly reduced in aged control mitochondria compared with young controls (P = 4.1 × 10^-5^; Figure 6F). In contrast, butyrate-treated aged hearts exhibited significantly higher State 3 OCR than those from untreated aged hearts (P = 1.4 × 10^-4^), indicating improved respiratory function. Collectively, these results demonstrate that late-life sodium butyrate supplementation attenuates age-associated alterations in cardiac mitochondrial ultrastructure and improves respiratory function.

### Butyrate supplementation enhances cardiac autophagy and mitophagy

Because the preceding analyses identified treatment-associated changes in autophagy-related proteins, autophagy-associated transcripts, and mitochondrial integrity, we next tested whether late-life butyrate supplementation was accompanied by enhanced cardiac autophagic activity. To examine these responses before advanced cardiac aging, we performed biochemical and reporter-based analyses in mice treated with sodium butyrate for three months beginning at 10 months of age and assessed at 13 months of age, an age-associated with emerging changes in cardiac structure and diastolic physiology^42^. Because steady-state autophagy markers alone cannot distinguish increased autophagosome formation from impaired lysosomal clearance^5, 21–23^, we assessed autophagic flux using chloroquine-mediated lysosomal inhibition and tandem fluorescent mRFP-GFP-LC3 reporter imaging^22, 57^, and mitophagy using mitochondria-targeted Keima^22, 58^.

We first examined LC3 processing in the presence or absence of chloroquine (CQ), a lysosomal inhibitor that blocks autophagosome degradation and thereby permits assessment of flux by measuring LC3-II accumulation. Representative immunoblotting demonstrated increased LC3-II abundance in butyrate-treated hearts relative to controls, both under basal conditions and following CQ treatment (Figure 7A). Quantification of LC3-II normalized to GAPDH showed significantly higher LC3-II levels in butyrate-treated samples than in control samples in the absence of CQ (P = 5.2 × 10^-5^) and in the presence of CQ (P = 3.5 × 10^-5^; Figure 7B). The further increase in LC3-II with CQ in butyrate-treated samples is consistent with enhanced autophagic flux rather than simple blockade of degradation, indicating that butyrate supplementation is associated with increased autophagosome turnover in cardiac tissue.

**Figure 7.**
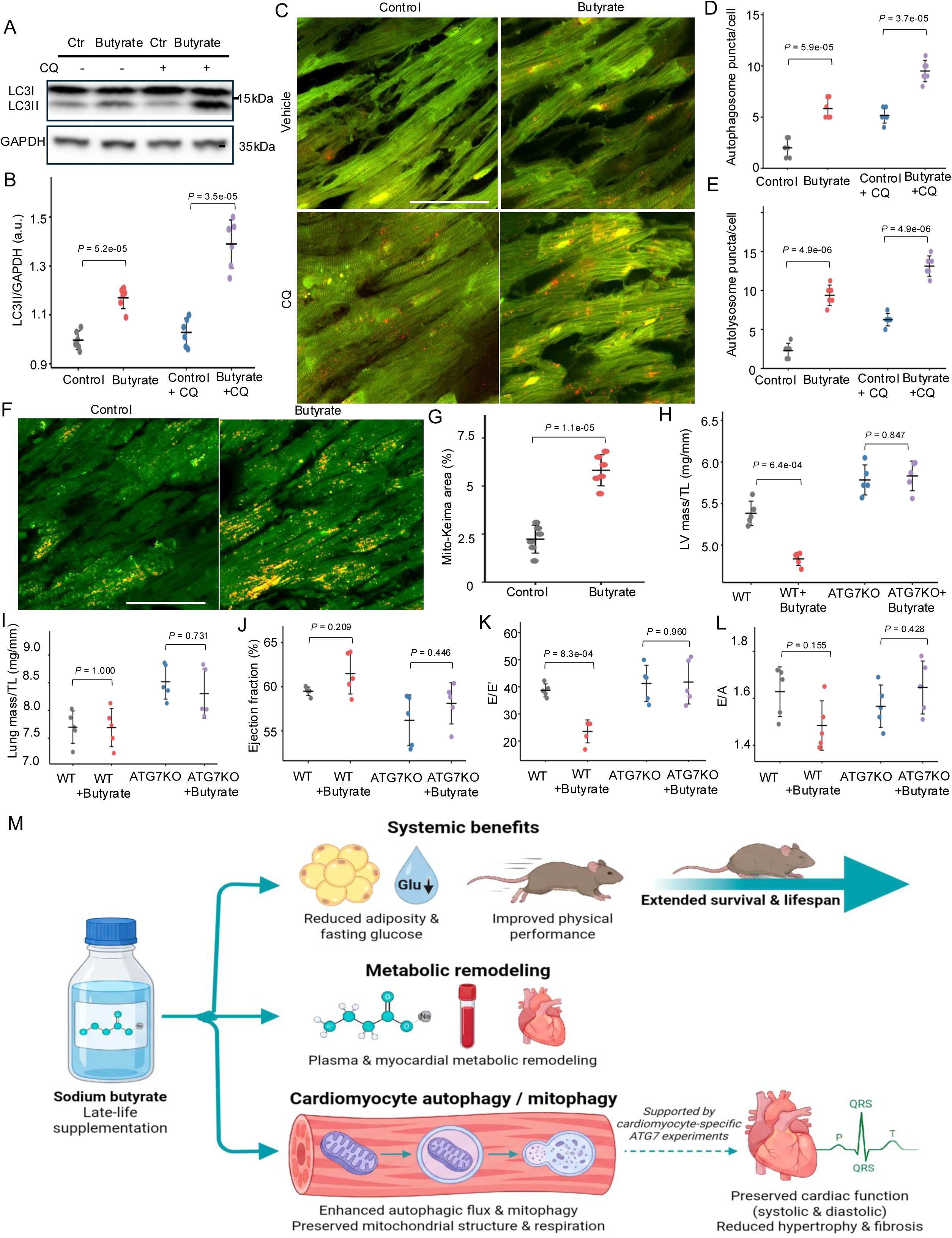
Late-life sodium butyrate supplementation enhances cardiac autophagic mitophagy. (A) Representative immunoblot of LC3-I and LC3-II in cardiac lysates from control and butyrate-treated mice, with or without chloroquine (CQ). GAPDH served as a loading control. (B) Quantification of LC3-II normalized to GAPDH, expressed in arbitrary units (a.u.). (C) Representative tandem fluorescent mRFP-GFP-LC3 (Tf-LC3) images of control and butyrate-treated myocardium under vehicle and CQ conditions. Autophagosomes appear as GFP⁺/RFP⁺ puncta and autolysosomes as predominantly RFP⁺ puncta. Scale bar, 50 μm. (D and E) Quantification of autophagosome (D) and autolysosome (E) puncta per cell. (F) Representative mito-Keima images of control and butyrate-treated myocardium. Acid-shifted fluorescence indicates mitochondrial material within acidic compartments. Scale bar, 50 μm. (G) Quantification of mito-Keima-positive area (%). For panels A–G, mice received butyrate supplementation for three months beginning at approximately 10 months of age and were assessed at 13 months. (H–L) Cardiac phenotyping of wild-type (WT) and cardiomyocyte-specific ATG7-deficient (ATG7KO) mice subjected to high-fat diet (HFD) plus L-NAME for four weeks, with or without butyrate supplementation. Treatment began at six weeks of age, and assessments were performed at approximately 10 weeks. (H) Left ventricular (LV) mass normalized to tibial length (TL). (I) Lung mass normalized to TL. (J) Ejection fraction (%). (K) Ratio of early transmitral filling velocity to early diastolic mitral annular velocity (E/E′). (L) Ratio of early to late transmitral filling velocities (E/A). Individual points represent biologically independent mice, and data are presented as mean ± SD. Pairwise comparisons in the ATG7 experiments were performed using two-sided, independent, equal-variance Student’s t-tests, followed by BH correction.

To visualize autophagic structures directly, we next used tandem fluorescent mRFP-GFP-LC3 (Tf-LC3), in which autophagosomes are identified as GFP-positive/RFP-positive puncta and autolysosomes as RFP-positive/GFP-negative puncta; GFP fluorescence is quenched in the acidic lysosomal environment thus changing the ratio^59, 60^. Representative images revealed increased punctate LC3 structures in butyrate-treated myocardium compared with controls under vehicle conditions and after CQ treatment (Figure 7C and Supplementary Figure 8). Quantification showed that butyrate treatment significantly increased autophagosome puncta per cell under basal conditions (P = 5.9 × 10^-5^) and after CQ administration (P = 3.7 × 10^-5^; Figure 7D). Autolysosome puncta per cell were likewise significantly increased by butyrate under vehicle (P = 4.9 × 10^-6^) and CQ-treated conditions (P = 4.9 × 10^-6^; Figure 7E), indicating that butyrate enhances autophagic flux in the heart rather than merely causing accumulation of undegraded vesicles.

Because mitochondrial structural and functional preservation was a prominent feature of the butyrate response, we next evaluated mitophagy using the pH-sensitive mitochondrial reporter mito-Keima. In this assay, increased acid-shifted mito-Keima signal reflects mitochondrial delivery to acidic lysosomal compartments and is therefore consistent with mitophagy-associated processing^57, 58, 60^. Representative images demonstrated a greater mito-Keima signal in butyrate-treated myocardium than in controls (Figure 7F). Quantification confirmed a significant increase in mito-Keima-positive area in butyrate-treated samples (P = 1.1 × 10^-5^; Figure 7G). These findings indicate that butyrate supplementation is associated with increased mitochondrial delivery to lysosomal compartments, consistent with enhanced mitophagy.

Previous studies have established the importance of ATG7-dependent autophagy and mitophagy in maintaining cardiac function^22, 61^. Notably, spermidine supplementation was shown to attenuate age-associated cardiac hypertrophy and diastolic dysfunction while enhancing autophagy, mitophagy, and mitochondrial respiration; these cardioprotective effects were impaired in mice with cardiomyocyte-specific deletion of Atg5^62^. To investigate whether core autophagy machinery similarly contributes to the cardiac responses to butyrate, we used mice with cardiomyocyte-specific ATG7 deficiency (Atg7fl/fl; α-MHC-Cre⁺, ATG7KO)^22^. Because prolonged cardiomyocyte autophagy deficiency can lead to progressive cardiac abnormalities, we examined these mice at an early age in a separate two-hit model of heart failure with preserved ejection fraction (HFpEF)^38, 63, 64^. This model combines high-fat diet (HFD) with nitric oxide synthase inhibition using Nω-nitro-L-arginine methyl ester (L-NAME) and reproduces several features relevant to cardiac aging, including ventricular hypertrophy, myocardial fibrosis, diastolic dysfunction, metabolic remodeling, and mitochondrial quality-control abnormalities^38, 63, 64^. Previous studies demonstrated that HFD plus L-NAME induces cardiac hypertrophy and diastolic dysfunction while maintaining relatively preserved ejection fraction after five weeks of treatment^63^, with similar cardiometabolic phenotypes reported after four and seven weeks in C57BL/6J mice^38^. We initiated HFD plus L-NAME exposure at six weeks of age and performed echocardiography and terminal assessments after four weeks, at approximately 10 weeks of age. This design allowed us to evaluate selected cardiac responses to butyrate under defined cardiometabolic stress while limiting the duration of cardiomyocyte ATG7 deficiency.

In WT mice, butyrate supplementation significantly reduced LV mass normalized to tibial length (P = 6.4 × 10^-4^; Figure 7H) and lowered the E/E′ ratio (P = 8.3 × 10^-4^; Figure 7K), consistent with attenuation of hypertrophic remodeling and improved diastolic indices. These treatment-associated differences were not statistically detected in ATG7KO mice (LV mass/TL, P = 0.847; E/E′, P = 0.960). Lung mass normalized to tibial length, ejection fraction, and E/A showed no significant treatment-associated differences within either genotype (Figure 7I, J, and L). Thus, the selective responses observed in WT mice, but not detected in ATG7KO mice, suggest a potential contribution of ATG7-related processes to butyrate-associated cardiac responses under cardiometabolic stress.

Collectively, late-life butyrate supplementation was associated with extended survival, improved systemic metabolic and physical function, and attenuation of cardiac aging (Figure 7M). Integrated multi-omics, mitochondrial analyses, and complementary autophagy and mitophagy experiments revealed metabolic remodeling, improved mitochondrial structure and respiration, and increased intracellular quality-control-associated activity. These findings highlight mitochondrial maintenance and autophagy-related pathways as potential contributors to the cardiac benefits of butyrate during aging.

## Discussion

In this study, we demonstrate that sodium butyrate supplementation initiated at 18 months of age extends subsequent survival in male and female C57BL/6J mice and attenuates multiple features of systemic and cardiac aging. Treatment reduced adiposity and fasting glucose, altered respiratory metabolism, and improved physical performance. In the heart, butyrate attenuated hypertrophic and fibrotic remodeling and improved systolic and diastolic functional indices. Integrated multi-omics analyses revealed distinct circulating and myocardial metabolic responses, accompanied by remodeling of ECM, RNA-processing, and autophagy-associated programs. These changes were associated with improved mitochondrial ultrastructure and respiratory function, while complementary reporter and genetic experiments implicated intracellular quality-control pathways as potential contributors to the cardiac response.

A central finding is that metabolic intervention initiated after substantial age-associated changes have accumulated can modify both survival and physiological function. The survival benefits observed in independent male and female cohorts were accompanied by improvements in adiposity, glucose regulation, and physical performance, suggesting a broader response beyond any single physiological endpoint. Although metabolic and nutrient-sensing interventions have shown promise in experimental aging^1, 2, 6^, our findings extend this framework to prolonged supplementation initiated during later life in a relatively long-lived mouse strain^30^. These convergent outcomes implicate further investigation of butyrate as a modulator of late-life physiological resilience.

The cardiac findings are particularly relevant because age-associated myocardial remodeling can progress without overt cardiovascular disease. Cardiac aging involves hypertrophy, fibrosis, mitochondrial dysfunction, and impaired relaxation, with systolic dysfunction emerging in some advanced-age cohorts^3, 4, 6, 42, 43^. Medrano et al. reported reduced EF in C57BL/6J mice at 29–31 months^42^, while Signore et al. documented reduced E/A, prolonged IVRT, and impaired ventricular relaxation beginning at approximately 25 months, followed by a reduction in EF at approximately 30 months^44^. More recently, Walter et al. reported age-associated cardiac hypertrophy and reduced EF in 24-month-old C57B6J substrain mice^65^. These studies provide context for the lower EF and E/A, higher E/E′, and prolonged IVRT observed in our 28-month-old controls. Butyrate supplementation attenuated LV hypertrophy and improved these functional indices in both sexes, suggesting that structural and functional cardiac aging remain responsive to intervention at an advanced age.

Our metabolomic findings extend previous investigations of diet–microbiota–host interactions into cardiac aging. Earlier studies from our group demonstrated individualized microbial and host responses to dietary fiber^12^, epigenetic regulation by butyrate and propionate^13^, and microbiome-associated metabolic signatures linked to chronic systolic heart failure^14^. Here, prolonged butyrate supplementation selectively remodeled circulating central-carbon intermediates, SCFAs, nucleotide-related metabolites, and host–microbial co-metabolites. Several metabolites elevated during aging, including citrate, succinate, α-ketoglutarate, acetate, and hydroxybutyrates, decreased following treatment, whereas others, notably IPA, increased further. IPA is of particular interest because previous studies reported its protective effects on diastolic function and cardiac metabolism in experimental HFpEF^38^ and implicated macrophage autophagy in its association with reduced aging-related myocardial fibrosis^39^. These findings suggest broader metabolic remodeling involving potentially cardioprotective metabolites.

Importantly, circulating metabolic changes did not uniformly reflect myocardial responses. Despite the observation that butyrate substantially elevated in plasma, myocardial abundance changed only modestly. Citrate and α-ketoglutarate decreased in plasma but increased in myocardium, whereas succinate decreased in both compartments. Particularly striking were trigonelline, hippuric acid, and 4-GBA, which increased in plasma but decreased in myocardium during aging, with the opposite responses following supplementation. These reciprocal patterns highlight compartment-dependent metabolic regulation and demonstrate that circulating metabolite abundance does not necessarily represent myocardial metabolic status^19, 20^. Trigonelline is of particular interest because it has been linked to NAD⁺ metabolism and muscle function during aging^45^, and its metabolic effects have also been investigated in experimental HFpEF^46^. Myocardial lipidomics similarly revealed selective changes in free fatty acids, lysophospholipids, and complex lipid species, potentially relevant to mitochondrial membrane organization and cellular quality control, but their subcellular localization and functional contributions remain to be determined.

Cardiac proteomics and histology provided complementary evidence of attenuated myocardial remodeling. Upon butyrate supplementation, ECM-associated pathways, including laminin interactions, collagen degradation, and ECM proteoglycans, showed enrichment patterns opposing those associated with aging, accompanied by reduced collagen-positive myocardial area. These findings support attenuation of fibrotic remodeling with supplementation, which may contribute to improved ventricular relaxation^4, 6, 43^. Proteomics also revealed opposing age-and treatment-associated enrichment of RNA-metabolism pathways and changes in RNA-processing proteins, motivating full-length RNA sequencing. Transcriptomic analysis identified selective remodeling of inflammatory, lipid-metabolism, ECM-associated, and autophagy-related programs, including transcript-level changes not apparent from parent-gene expression. Notably, Wipi1-202 decreased during aging and increased following supplementation despite no significant difference in total Wipi1 expression. WIPI1 participates in phosphoinositide-dependent autophagic membrane formation and autophagosome biogenesis^47, 48^ and has been implicated in cardiac dysfunction and mitochondrial quality control^50, 51^. These findings nominate Wipi1-202 as a potentially relevant transcript within the butyrate response.

Mitochondrial preservation represents another important feature of the cardiac response. Aging is associated with enlarged mitochondrial profiles, cristae disruption, and impaired oxidative phosphorylation^3, 52–56^. Consistent with these abnormalities, aged control hearts exhibited increased mitochondrial cross-sectional area and reduced cristae abundance and organization. Butyrate-treated hearts showed reduced mitochondrial aging profiles and better-preserved cristae architecture, accompanied by higher substrate-supported baseline and ADP-stimulated State 3 respiration. These findings indicate that mitochondrial ultrastructure and respiratory function remain responsive to supplementation initiated late in life. Together with the myocardial metabolic and molecular changes, they highlight mitochondrial maintenance as a potential component of butyrate-associated cardiac protection.

The association between autophagy-related remodeling and mitochondrial preservation provides a potential biological link between these findings. Beyond changes in autophagy-associated proteins and transcripts, LC3-II measurements following lysosomal inhibition, tandem fluorescent LC3 reporter imaging, and mito-Keima assays provided complementary evidence of altered autophagic activity and increased mitochondrial delivery to acidic compartments following butyrate supplementation. These experiments were conducted in separate, younger cohorts and therefore do not directly establish that enhanced autophagy mediates mitochondrial preservation in the 28-month-old hearts. Nevertheless, the convergence of molecular, ultrastructural, respiratory, and reporter findings supports intracellular quality control as a candidate pathway contributing to the cardiac response.

Previous dietary-intervention studies provide a relevant mechanistic precedent. Eisenberg et al. demonstrated that spermidine supplementation extended lifespan, improved cardiac hypertrophy and diastolic function, and enhanced autophagy, mitophagy, and mitochondrial respiration; cardioprotection was impaired following cardiomyocyte-specific deletion of Atg5^62^. Similarly, studies of HFD plus L-NAME-induced HFpEF have implicated impaired mitophagy and mitochondrial respiration in cardiometabolic dysfunction^63, 64^. In our separate two-hit HFpEF model, butyrate reduced LV mass and E/E′ in WT mice, whereas corresponding improvements were not statistically detected in cardiomyocyte-specific ATG7-deficient mice. These observations are consistent with a potential contribution of ATG7-dependent processes to selected cardiac responses. However, a significant effect in WT mice and a nonsignificant effect in ATG7-deficient mice do not independently establish genotype dependence. Formal genotype-by-treatment interaction analysis and genetic studies in naturally aged mice will be needed to determine whether ATG7-dependent autophagy is required for the benefits observed during advanced aging.

Our findings also extend previous experimental studies showing that butyrate attenuates cardiac dysfunction, hypertrophy, and fibrosis in diet-induced metabolic disease^17^ and affects cardiovascular function under other experimental stress conditions^66^. Unlike these disease-focused models, the present study examines prolonged supplementation initiated during natural aging. Emerging clinical evidence provides additional translational context. A randomized, placebo-controlled trial reported that oral sodium butyrate combined with moderate dietary energy restriction improved weight-related outcomes and selected metabolic measures in adults with overweight, obesity, or type 2 diabetes^18^, whereas earlier work demonstrated differences in metabolic responsiveness between lean individuals and those with metabolic syndrome^67^. Although these studies support the potential for oral butyrate to influence human metabolism, its effects on cardiac aging remain unknown. Differences in treatment duration, exposure, and metabolic context will need to be addressed before translating the present findings to older human populations.

Collectively, our findings demonstrate that late-life butyrate supplementation is associated with longer survival and attenuation of multiple systemic and cardiac aging phenotypes. Distinct circulating and myocardial metabolic responses, remodeling of ECM and RNA-processing programs, preservation of mitochondrial structure and respiration, and enhanced autophagy-and mitophagy-associated activity identify metabolic regulation and intracellular quality control as promising pathways for understanding and potentially modifying cardiac aging.

### Limitations of the study

Several considerations should be acknowledged. The survival, physiological, multi-omics, and mechanistic findings were obtained from complementary experimental cohorts, providing convergent evidence across multiple levels of biological organization but not establishing a single causal pathway linking butyrate supplementation to improved survival and cardiac function. In addition, circulating and myocardial metabolomics, lipidomics, and bulk tissue transcriptomics identify treatment-associated molecular changes without resolving metabolic flux, cell-type-specific responses, or the functional contributions of individual metabolites and RNA isoforms. Although mitochondrial imaging, respiration assays, and autophagy reporters provide complementary evidence of improved mitochondrial integrity and increased quality-control-associated activity, additional targeted studies will be needed to define the underlying molecular mechanisms.

The ATG7-deficiency experiments were conducted in a separate two-hit HFpEF model rather than in naturally aged mice, and the observed within-genotype treatment differences require formal genotype-by-treatment interaction testing to establish ATG7 dependence. Furthermore, the mechanisms underlying the systemic and cardiac responses may involve both direct myocardial effects and indirect metabolic or vascular influences. Future studies combining targeted genetic perturbation, cell-type-resolved analyses, and metabolic tracing will help determine how butyrate-associated systemic remodeling contributes to cardiac protection and whether these effects extend to human cardiovascular aging.

## RESOURCE AVAILABILITY

### Lead contact

Further information and requests for resources and reagents should be directed to the lead contact, Michael P. Snyder.

### Materials availability

Reagents and mouse lines described in this study will be made available upon reasonable request to the lead contact, subject to availability and applicable institutional, biosafety, and material transfer requirements.

### Data and code availability

Full-length RNA-sequencing data will be deposited in the Gene Expression Omnibus. Proteomics, metabolomics, lipidomics, and processed physiological data will be deposited in appropriate public repositories or provided as supplementary source data, with accession numbers and links supplied upon availability. Additional information required to reanalyze the data will be available from the lead contact upon reasonable request.

## Acknowledgments

We thank Junichi Sadoshima (Rutgers New Jersey Medical School) for providing the tandem fluorescent LC3 (tf-LC3) and mito-Keima reporter mice. We also thank Lucia Ramirez and proteomics team members of the Snyder laboratory for their assistance and discussions. We acknowledge the Stanford University Mass Spectrometry (SUMS) facility, Stanford Cell Sciences Imaging Facility (CSIF), and Behavioral and Functional Neuroscience Laboratory (BFNL) for their technical support and access to instrumentation. We thank the Stanford Diabetes Research Center (SDRC), supported by NIH grant P30DK116074, for access to the CLAMS/Oxymax metabolic phenotyping system. We also acknowledge the Stanford imaging facilities for access to the Vevo F2 ultrasound imaging system and 11.7-T MRI system. This work was supported gifts from Embrace Innovations Research Foundation and the Anu and BV Jagadeesh Family Foundation. M.T. was supported by an NIH National Research Service Award (NRSA) postdoctoral fellowship (F32HL182300) and an NIH institutional training grant (T32DK007217).

## Author contributions

Conceptualization, M.T. and M.P.S.; Methodology, M.T., L.J., R.J., F.C., F.W., and T.C.; Investigation, M.T., L.J., R.J., F.W., T.C.; Formal Analysis, M.T.; Data Curation, M.T.; Visualization, M.T. and L.M.; Supervision, M.P.S.; Funding Acquisition, M.T. and M.P.S.; Writing – Original Draft, M.T. and M.P.S.; Writing – Review & Editing, all authors.

## Delaration of interests

The authors declare no competing interests.

## Delaration of generative AI and AI-assisted technologies in the writing process

ChatGPT was used to improve grammar and readability; authors reviewed all content and take full responsibility for the publication.

## STAR METHODS

### KEY RESOURCES TABLE

A complete Key Resources Table will accompany the final manuscript and identify the sources, catalog numbers, and resource identifiers for mouse strains, antibodies, chemicals and software used in the study. Key resources include C57BL/6J mice from The Jackson Laboratory; cardiomyocyte-specific tandem fluorescent mRFP-GFP-LC3 and mito-Keima reporter mice; cardiomyocyte-specific ATG7-deficient mice; sodium butyrate; chloroquine; LC3 and GAPDH antibodies; isotope-labeled metabolite standards; lipid internal standards; and the sequencing, mass-spectrometry, microscopy, echocardiography, and metabolic phenotyping platforms described below.

### EXPERIMENTAL MODEL AND STUDY PARTICIPANT DETAILS

All animal procedures were approved by the Stanford University Institutional Animal Care and Use Committee and conducted in accordance with the NIH Guide for the Care and Use of Laboratory Animals. Mice were maintained under specific pathogen-free conditions on a 12-h light/dark cycle with standard chow and water available ad libitum, except during specified fasting procedures.

Male and female C57BL/6J mice were obtained from The Jackson Laboratory at approximately 18 months of age and assigned to control or sodium butyrate groups. Sodium butyrate was administered in drinking water at 10 g/L (1% w/v), while controls received regular drinking water. Treatment continued from 18 to 28 months for terminal phenotyping or until death for survival analyses. Young-reference mice were 4 months old.

Separate cohorts were used for survival, metabolic and physical assessments, cardiac phenotyping, multi-omics, mitochondrial analyses, and autophagy experiments, as indicated in the figure legends. Individual mice served as biological replicates. Autophagy reporter experiments were conducted in separate cohorts treated from approximately 10 to 13 months of age, whereas ATG7-deficiency experiments were performed in a distinct two-hit HFpEF model.

Cardiac autophagy and mitophagy were evaluated using cardiomyocyte-specific tandem fluorescent mRFP-GFP-LC3 (Tf-LC3) and mitochondria-targeted Keima reporter mice, respectively^22^. Cardiomyocyte-specific ATG7-deficient mice (Atg7fl/fl; α-MHC-Cre⁺, ATG7KO) were generated using floxed Atg7 mice and an α-myosin heavy chain Cre driver, as described previously^22^.

To investigate ATG7-dependent cardiac responses to butyrate, wild-type (WT) and ATG7KO mice were subjected to a two-hit heart failure with preserved ejection fraction (HFpEF) model combining high-fat diet (HFD) and the nitric oxide synthase inhibitor Nω-nitro-L-arginine methyl ester (L-NAME), as described previously^63^. Beginning at six weeks of age, mice received HFD plus L-NAME with or without butyrate supplementation for four weeks. Echocardiography and terminal tissue collection were performed at approximately 10 weeks of age. LV and lung masses were normalized to tibial length, and cardiac function was evaluated using ejection fraction, E/E′, and E/A. Individual mice served as biological replicates.

### METHOD DETAILS

#### Survival analysis

Independent survival experiments were conducted in male and female C57BL/6J mice, with 24 control and 24 sodium butyrate-treated animals per sex. Mice entered the experiments at approximately 18 months of age, when supplementation was initiated. Survival was monitored until death, and age at death was recorded in days. Kaplan–Meier survival curves were generated using the R package survival. Survival distributions were compared using a two-sided log-rank test, and median survival was estimated from the Kaplan–Meier curves. Dashed horizontal and vertical reference lines in the survival plots indicate 50% survival and estimated median survival times. Because supplementation began at 18 months of age, the analyses describe survival among animals that had survived to treatment initiation. Chronological age at death was used for plotting, rather than treating animals as having received the intervention from birth.

#### Plasma butyrate quantification

Plasma butyrate concentrations were determined using a targeted 3-nitrophenylhydrazine (3-NPH) derivatization liquid chromatography–tandem mass spectrometry (LC–MS/MS) assay. Plasma samples were analyzed using an Agilent 6495 triple-quadrupole mass spectrometer. A ^13^C₄-labeled butyrate internal standard was added to samples, and concentrations were calculated using a calibration series containing known butyrate concentrations. Three biologically independent aged male mice per group were analyzed. Plasma butyrate concentrations were expressed in μmol/L. This calibrated assay was distinguished from the separate relative-abundance measurement of butyrate included in the broader targeted organic acid dataset.

#### Longitudinal body weight and fasting blood glucose

Body weight was recorded monthly from 18 to 28 months of age in male and female control and butyrate-treated cohorts, with six independent mice per group. Fasting blood glucose was measured at 18, 21, 24, and 28 months of age, with six mice per group at each assessment. Male and female datasets were analyzed separately. The duration of fasting, sampling site, and glucose measurement device should be specified from the experimental records. Body-weight measurements collected during indirect calorimetry were analyzed separately from the longitudinal body-weight dataset.

#### Abdominal adiposity assessment

At terminal assessment, epididymal and perirenal adipose tissue depots were dissected and weighed. The combined mass of the dissected adipose tissue was recorded in grams. The abdominal fat index was calculated as: Abdominal fat index (%) = [(epididymal fat mass + perirenal fat mass) / total body weight at dissection] × 100. Absolute combined epididymal and perirenal fat mass and the abdominal fat index were analyzed as separate endpoints. These measurements were obtained directly from dissected adipose tissue.

#### Anatomical magnetic resonance imaging

Representative anatomical MRI were acquired at 11.7 T using ParaVision 360 software, version 2.0 PL1. Mice were maintained under approximately 1%–2% isoflurane anesthesia, adjusted to maintain a respiratory rate of approximately 70–80 breaths/min. Physiological monitoring was performed using PC-SAM. Localizer imaging and radiofrequency tuning and matching preceded acquisition of coronal T2-weighted rapid acquisition with relaxation enhancement (RARE) images. Images were exported in DICOM format. The images were used for qualitative anatomical visualization and were not used for quantitative abdominal fat measurements.

#### Indirect calorimetry and metabolic phenotyping

Indirect calorimetry was performed at the Stanford Metabolic Core using a Comprehensive Lab Animal Monitoring System (CLAMS)-HC Oxymax system (Columbus Instruments). Approximately 20-month-old male control and butyrate-treated mice were individually housed in metabolic monitoring chambers for seven days, with four biologically independent mice per group. The first 48 h were designated as acclimation, and measurements collected during days 3–7 were used for analysis. Oxygen consumption (VO₂), carbon dioxide production (VCO₂), respiratory exchange ratio (RER), food intake, and calculated heat production were recorded. RER was calculated as VCO₂ divided by VO₂. Measurements were summarized separately for the light and dark phases. Repeated instrument measurements were averaged within each mouse and phase before statistical analysis, so that each animal contributed one summary observation per phase and endpoint. Body weight was analyzed once per animal for the monitoring session, rather than separately for the light and dark phases. Daily water intake was also assessed in the metabolic phenotyping cohort. Heat production was reported normalized to body weight in kcal/kg/h. Because body-weight-normalized energy expenditure may be influenced by variation in body size and composition, these measurements were interpreted descriptively rather than as evidence of energy expenditure independent of body composition^32, 33^. The male indirect-calorimetry statistical comparison family comprised 11 unique comparisons: light-and dark-phase comparisons for food intake, heat production, VCO₂, VO₂, and RER, together with a single session-level body-weight comparison.

#### Open-field locomotor activity

Spontaneous locomotor activity was assessed using an open-field test in young, aged control, and aged butyrate-treated male mice, with five independent mice per group. Mice were individually placed in the center of an opaque 50 × 50 cm arena under approximately 30-lux illumination. Activity was recorded for 60 min and tracked using SMART software, version 3.0 (Panlab). Total distance traveled and vertical activity were analyzed as separate endpoints. Total distance was expressed in meters following conversion from the original tracking-export units. Between animals, the arena was cleaned with disinfectant and allowed to stand for approximately 2 min.

#### Grip-strength assessment

Muscle strength was evaluated using a grip-strength meter in young, aged control, and aged butyrate-treated mice. Two-paw and four-paw grip-strength measurements were obtained from the same animals. Mice were allowed to grasp the appropriate bar or grid and were gently pulled backward until grip release. The recorded force was expressed in newtons. Six biologically independent mice were analyzed per group for each sex. The two grip-strength measurements were treated as distinct outcomes obtained from the same biological subjects.

#### Accelerating rotarod test

Motor coordination and performance were assessed using an accelerating rotarod apparatus (Panlab) equipped with a 30-mm-diameter rotating rod. Rotational speed increased from 5 to 40 rpm over 5 min. Two trials were conducted daily, separated by approximately 2 h, for five consecutive days. Each trial ended when the mouse fell from the rod or reached the maximum observation period. Latency to fall during the final trial was used for quantitative comparisons and expressed in seconds. Six biologically independent mice were included per group for each sex. Young reference mice were 4 months old, and aged mice were assessed at 28 months.

#### Organ collection and tissue mass measurements

At terminal assessment, hearts and lungs were collected and weighed. Left ventricular tissue mass and lung mass were normalized to tibial length to account for variation in animal size. Normalized organ mass was calculated as organ mass (mg) divided by tibial length (mm). Cardiac and pulmonary mass measurements were compared among young, aged control, and aged butyrate-treated mice, with six mice per group for the main cardiac phenotyping analyses.

#### Echocardiography

Cardiac structure and function were assessed using a VisualSonics Vevo F2 ultrasound imaging system equipped with a UHF46x transducer. Mice underwent transthoracic echocardiography under controlled anesthesia, and body temperature was maintained at approximately 37°C using a temperature-controlled heating pad. Standard imaging planes, M-mode recordings, pulsed-wave Doppler recordings, and tissue Doppler images were acquired and stored for offline analysis.The parasternal long-axis view was used to evaluate LV dimensions and wall motion. Representative M-mode recordings were used to illustrate ventricular wall motion and chamber geometry. Transmitral pulsed-wave Doppler and mitral annular tissue Doppler recordings were obtained from an apical long-axis view. Early transmitral filling velocity (E) and late transmitral filling velocity associated with atrial contraction (A) were measured, and the E/A ratio was calculated. Early diastolic mitral annular velocity (E′) was measured using tissue Doppler imaging. The E/E′ ratio was calculated using consistent velocity units. Isovolumic relaxation time (IVRT) was measured as the interval between aortic valve closure and mitral valve opening. Left ventricular ejection fraction was calculated from echocardiographic measurements using the imaging software. E/A, E/E′, and IVRT were interpreted collectively as indices of diastolic function, recognizing that these measurements are influenced by heart rate, loading conditions, and anesthesia^7, 8^. Six biologically independent mice were analyzed per group for each sex in the main cardiac phenotyping experiments. Image acquisition and quantitative analyses were performed with blinding to treatment assignment where documented.

#### Plasma metabolomics and lipidomics

Sample preparation and extraction Plasma samples from young, aged control, and aged butyrate-treated male mice were analyzed using targeted 3-NPH organic acid profiling, broader metabolomic profiling, and lipidomic profiling, with three biologically independent mice per group for each dataset. A combined extraction procedure was used to separate lipid-containing organic fractions and polar-metabolite-containing fractions. For plasma extraction, 100 μL of plasma was transferred to a 2-mL microcentrifuge tube. Ice-cold methanol (500 μL) was added, followed by 10 μL of EquiSPLASH lipid internal-standard mixture. Samples were vortexed for approximately 20 s, and 1,000 μL ice-cold methyl tert-butyl ether (MTBE) was added. Following vortexing, 250 μL ice-cold water was added and samples were vortexed for 1 min. Samples were incubated at −20°C for 10–60 min and centrifuged at 15,000 × g for 15 min at 4°C. The upper organic phase was collected for lipidomic analysis. The lipid-containing fraction was dried under nitrogen and reconstituted in acetonitrile/dichloromethane (93:7, v/v). For polar-metabolite extraction, the remaining material was processed with 500 μL ice-cold acetonitrile/methanol (1:1, v/v). CIL QReSS isotope-labeled metabolite internal standards were added at 10 μL per sample. Samples were incubated at −20°C for 1 h to overnight and centrifuged at 10,000 × g for 10 min at 8°C. The recovered supernatant was used for broader metabolomic profiling and targeted 3-NPH analysis.

#### Targeted 3-NPH derivatization and LC–MS/MS

A 50-μL aliquot of polar-metabolite-containing supernatant was removed before evaporation and subjected to 3-NPH derivatization. Fresh 3-NPH and 1-ethyl-3-(3-dimethylaminopropyl)carbodiimide (EDC) solutions were prepared for each derivatization batch. The 3-NPH solution contained 200 mM 3-NPH in water/acetonitrile (1:1, v/v). The EDC solution contained 120 mM EDC in water/acetonitrile/pyridine (47:47:6, v/v/v). Following addition of the assay-specific internal standard, 20 μL of 3-NPH solution and 20 μL of EDC solution were added. Samples were mixed and incubated at 37°C for 45 min with shaking. The reaction was quenched with 20 μL of 0.1% formic acid in water/acetonitrile (80:20, v/v), followed by an additional 10-min incubation at 37°C. Derivatized samples were diluted 50-fold in water/acetonitrile (90:10, v/v) before analysis on an Agilent 6495 triple-quadrupole mass spectrometer. The targeted acquisition method used negative-ion multiple-reaction monitoring for the specified organic acids. Butyrate-d₃ was used as the internal standard for the relative-abundance butyrate measurement in the targeted organic acid dataset.

#### Broad plasma metabolomic profiling

The polar-metabolite fraction was prepared in water/methanol/acetonitrile (2:1:7, v/v/v) and analyzed using an Agilent 6496 mass spectrometry platform. Metabolite abundances were used for individual-feature comparisons, principal component analysis (PCA), visualization of selected metabolites, and classification of significantly altered features into biochemical categories. Metabolite abundances were reported as relative measurements unless otherwise specified. Row-standardized z scores were used for visualization in heatmaps.

#### Plasma lipidomic profiling

The lipid-containing MTBE fraction was analyzed using a SCIEX 7500+ mass spectrometry platform. EquiSPLASH internal standards were introduced during extraction. Selected lipid features included phosphatidylcholines, lysophosphatidylcholines, phosphatidylethanolamines, phosphatidylinositols, phosphatidylserines, hexosylceramides, diacylglycerols, and triacylglycerols. Lipid species were annotated according to the structural information supported by the analytical method and established lipid nomenclature^41^. Partial fatty-acyl annotations were not interpreted as determining exact double-bond locations or positional isomers.

#### Left ventricular metabolomics and lipidomics

Left ventricular tissue from young, aged control, and aged butyrate-treated male mice was analyzed using complementary metabolomic and lipidomic platforms, with three biologically independent mice per group. Approximately 40 mg of LV tissue was homogenized and processed using a combined extraction strategy adapted from the plasma workflow. Organic and polar fractions were separated for lipidomic and metabolomic analyses, respectively. EquiSPLASH lipid standards and CIL QReSS isotope-labeled metabolite standards were added during the corresponding extraction steps. Targeted 3-NPH analyses were performed using the Agilent 6495 platform, broader metabolomic profiling using the Agilent 6496 platform, and lipidomic profiling using the SCIEX 7500+ platform. Myocardial metabolite and lipid measurements were reported as relative abundance unless otherwise indicated. Treatment-associated metabolite changes were compared between plasma and LV tissue using log₂ fold changes calculated separately within each compartment. Because plasma and LV measurements represent distinct biological compartments, these comparisons were used to characterize similarities and differences in relative abundance responses rather than to infer metabolite transfer or flux between circulation and myocardium.

#### Left ventricular proteomics

LV tissue from young, aged control, and aged butyrate-treated male mice was subjected to data-independent acquisition (DIA) proteomic analysis, with three biologically independent mice per group. Peptides were analyzed using an Orbitrap Astral mass spectrometer. Raw mass spectrometry data were processed using DIA-NN in a library-free workflow. Peptide and precursor identification filtering used a 1% false-discovery rate (FDR). Protein differential-abundance analysis was conducted for aged control versus young, aged butyrate-treated versus aged control, and aged butyrate-treated versus young comparisons. Protein-level P values were adjusted using the Benjamini–Hochberg procedure. Differentially abundant proteins were defined using an adjusted FDR threshold below 0.05. Protein fold-change patterns were compared between aging-and treatment-associated contrasts to identify proteins with concordant or opposing abundance responses. Pathway enrichment analyses included Hallmark, Reactome, and Gene Ontology Biological Process collections. Normalized enrichment scores were used to determine enrichment direction, and significance was evaluated with multiple-testing correction across the corresponding tested pathway collection. Protein features associated with ECM organization, RNA metabolism, RNA splicing, protein folding, translation, mTORC1 signaling, autophagy, lysosomal function, and vesicle trafficking were examined.

#### Picrosirius red staining and cardiac fibrosis quantification

Myocardial fibrosis was assessed in aged control and aged butyrate-treated mice using picrosirius red (PSR) staining. Representative whole-section images and higher-magnification views of perivascular and interstitial regions were obtained to visualize collagen-containing structures. Collagen-positive areas were quantified using digital image analysis, including perivascular and interstitial collagen-positive regions within the analyzed myocardial tissue. Fibrosis area was calculated as: Fibrosis area (%) = (collagen-positive area / analyzed myocardial tissue area) × 100. Six biologically independent mice were analyzed per group. Multiple regions or images from the same mouse were summarized to obtain a biological measurement for each animal. The quantification procedure was based on segmentation of PSR-positive collagen-containing regions, with the final image-analysis criteria and thresholds specified from the experimental processing records^68^.

#### PacBio Kinnex full-length RNA sequencing

Full-length RNA sequencing was performed using LV tissue from young, aged control, and aged butyrate-treated male mice, with three biologically independent mice per group. RNA was extracted and processed using the PacBio Kinnex full-length RNA sequencing workflow. Full-length cDNA sequencing enabled characterization of transcript structures and quantification of individual transcripts and their corresponding parent genes. Transcript annotations were used to distinguish genes represented by single or multiple detected transcripts. Gene-level and transcript-level differential-expression analyses were performed for the indicated group comparisons. Transcripts were assigned to their parent genes to identify genes containing at least one significantly differentially expressed transcript even when the parent gene did not meet the gene-level significance threshold. PCA was used to characterize variation in transcript-expression profiles across experimental groups. Pathway enrichment analyses were conducted using ranked expression changes. Selected RNA-processing, inflammatory, lipid-metabolism, ECM-associated, autophagy, and mitophagy-related transcripts were examined. Transcript structures and abundances were evaluated for selected loci, including Ankrd23 and Wipi1. Relative transcript representation was calculated as the abundance of an individual detected transcript divided by the summed abundance of the corresponding detected transcripts from the same gene, multiplied by 100. Transcript-level differential expression and relative transcript usage were treated as distinct outcomes.

#### Transmission electron microscopy

LV tissue from young, aged control, and aged butyrate-treated male mice was processed for transmission electron microscopy (TEM) to assess mitochondrial morphology and cristae organization. Representative images were acquired at low and high magnification to visualize mitochondrial distribution and ultrastructural features within cardiomyocytes. Quantitative measurements included mitochondrial cross-sectional area, the number of discernible cristae per mitochondrial profile, and a semiquantitative cristae morphology score. Mitochondrial cross section area was measured from two-dimensional TEM sections and expressed in μm². This parameter was interpreted as profile area rather than mitochondrial volume. Cristae number was defined as the number of discernible cristae within each imaged mitochondrial profile, without normalization to mitochondrial area. Cristae morphology was evaluated using an ordinal scoring system, with higher scores corresponding to better-preserved internal membrane architecture. Three biologically independent mice were analyzed per group. The figure level analysis describes 50 mitochondrial profile measurements per mouse for each endpoint, subject to verification against the final source-data file. Mitochondrial profiles sampled from the same mouse were treated as nested observations.

#### Isolation of cardiac mitochondria and oxygen consumption measurement

Oxygen consumption measurement has been described previously^61, 64^. Briefly, cardiac mitochondria were isolated from young, aged control, and aged butyrate-treated mice. Fresh cardiac tissue was homogenized in ice-cold mitochondrial isolation buffer, and mitochondria were enriched by differential centrifugation. Mitochondrial protein concentrations were determined before extracellular flux analysis. Oxygen consumption rate (OCR) was measured using a Seahorse XF extracellular flux analyzer. Isolated mitochondria were suspended in mitochondrial assay solution supplemented with pyruvate (10 mM) and malate (2 mM) and loaded into the assay plate at a standardized protein concentration. Following centrifugation to facilitate mitochondrial attachment to the assay plate, mitochondria were equilibrated in assay medium containing the same respiratory substrates. Substrate-supported baseline OCR was measured in the presence of pyruvate and malate before ADP administration. ADP was subsequently injected to a final concentration of 5 mM, and ADP-stimulated State 3 OCR was measured. OCR was normalized to the amount of mitochondrial protein loaded per well and expressed as pmol O₂/min/μg mitochondrial protein. Technical replicate wells were averaged to obtain one value per mouse. Six biologically independent mice were analyzed per group.

#### LC3 immunoblotting and chloroquine-based autophagy assessment

Cardiac autophagy-associated activity was examined in a separate cohort of mice treated with sodium butyrate for three months beginning at approximately 10 months of age and assessing at 13 months of age. To assess LC3-II accumulation under conditions of lysosomal inhibition, control and butyrate-treated mice received chloroquine (CQ) or vehicle approximately 4 h before euthanasia. Cardiac lysates were prepared and subjected to immunoblotting for LC3-I, LC3-II, and GAPDH. Immunoreactive signals were quantified by densitometry, and LC3-II abundance was normalized to GAPDH. Treatment-associated differences in LC3-II abundance were evaluated in the absence and presence of CQ. Because steady-state LC3-II abundance may be affected by both formation and degradation, interpretation of autophagic flux was based on the response to lysosomal inhibition rather than basal LC3-II abundance alone^22^.

#### Tandem fluorescent LC3 reporter imaging

To investigate the effects of sodium butyrate on cardiac autophagy and mitophagy during middle age, mice were treated with sodium butyrate in drinking water for three months beginning at approximately 10 months of age and analyzed at 13 months. This age was selected to examine autophagy-associated responses during an earlier phase of cardiac aging, informed by previous observations of age-associated cardiac structural and diastolic changes in middle-aged mice^42^. Cardiac autophagic activity was evaluated using chloroquine-mediated LC3-II accumulation and cardiomyocyte-specific Tf-LC3 reporter mice. Mitophagy-associated activity was assessed using mice expressing mitochondria-targeted Keima in cardiomyocytes. These experiments were performed in age-defined cohorts separate from the 28-month natural-aging study. Cardiac autophagy-associated compartments were examined using mice with cardiomyocyte-specific expression of Tf-LC3. Reporter mice received sodium butyrate supplementation for three months beginning at approximately 10 months of age. Mice were treated with CQ or vehicle approximately 4 h before euthanasia, according to the experimental grouping. Cardiac tissue was imaged to visualize GFP and RFP signals separately and in merged images. Autophagosome-associated puncta were classified as dual-positive GFP/RFP structures, whereas autolysosome-associated puncta were classified as predominantly RFP-positive structures under conditions allowing quenching of GFP fluorescence in acidic compartments. Autophagosome-and autolysosome-associated puncta were quantified per cardiomyocyte and summarized for each biological sample. Interpretation of the reporter response considered the effects of lysosomal inhibition and CQ-associated changes in lysosomal acidification on fluorescence classification^57, 69^.

#### Mito-Keima measurement of mitophagy

Mitophagy was assessed using mice with cardiomyocyte-specific expression of mitochondria-targeted Keima. Mito-Keima is a pH-sensitive fluorescent reporter that undergoes an excitation shift when mitochondria are delivered to acidic lysosomal compartments^58^. Cardiac tissue was imaged using reporter-specific excitation and emission settings. Acid-shifted mito-Keima signal was used to identify mitochondrial material associated with acidic compartments. Mito-Keima-positive area was quantified and expressed as a percentage of the analyzed area, as displayed in Figure 7G. Measurements were summarized at the animal level for statistical comparison.

### QUANTIFICATION AND STATISTICAL ANALYSIS

Statistical analyses were performed using R. Data are presented as mean ± SD unless otherwise indicated. Individual mice were considered independent biological replicates, and sample sizes are specified in the corresponding figure legends. Between-group comparisons were performed using two-sided, unpaired Student’s t-tests assuming equal variances, followed by Benjamini–Hochberg (BH) correction for multiple comparisons where applicable. Survival distributions were analyzed using the Kaplan–Meier method and compared using the two-sided log-rank test. Gene-and transcript-level differential-expression analyses were performed using DESeq2, and differential protein abundance and pathway enrichment were assessed using the corresponding bioinformatics pipelines with multiple-testing correction. Genotype-by-treatment interactions were evaluated for the ATG7-deficient mouse experiments. A BH-adjusted P value or false discovery rate (FDR) < 0.05 was considered statistically significant where applicable. Specific statistical tests, sample sizes, and multiple-comparison procedures are described in the corresponding figure legends and Methods.

## ABBREVIATIONS

3-NPH: 3-Nitrophenylhydrazine
4-GBA: 4-Guanidinobutyric acid
ATG7: Autophagy-related protein
7 BH: Benjamini–Hochberg
CLAMS: Comprehensive Lab Animal Monitoring System
CQ: Chloroquine
DIA: Data-independent acquisition
E: Early transmitral filling velocity
E’: Early diastolic mitral annular velocity
ECM: Extracellular matrix
EF: Ejection fraction
FDR: False discovery rate
HFpEF: Heart failure with preserved ejection fraction
IPA: Indole-3-propionic acid
IVRT: Isovolumic relaxation time
LC3: Microtubule-associated protein 1 light chain 3
LV: Left ventricle/left ventricular
MRI: Magnetic resonance imaging
mt-Keima: Mitochondria-targeted Keima
NaBu: Sodium butyrate
OCR: Oxygen consumption rate
PCA: Principal component analysis
PSR: Picrosirius red
RBP: RNA-binding protein
RER: Respiratory exchange ratio
SCFA: Short-chain fatty acid
TEM: Transmission electron microscopy
Tf-LC3: Tandem fluorescent mRFP-GFP-LC3
TL: Tibial length
WT: Wild type

## Supplemental information

**Supplementary Figure 1.**
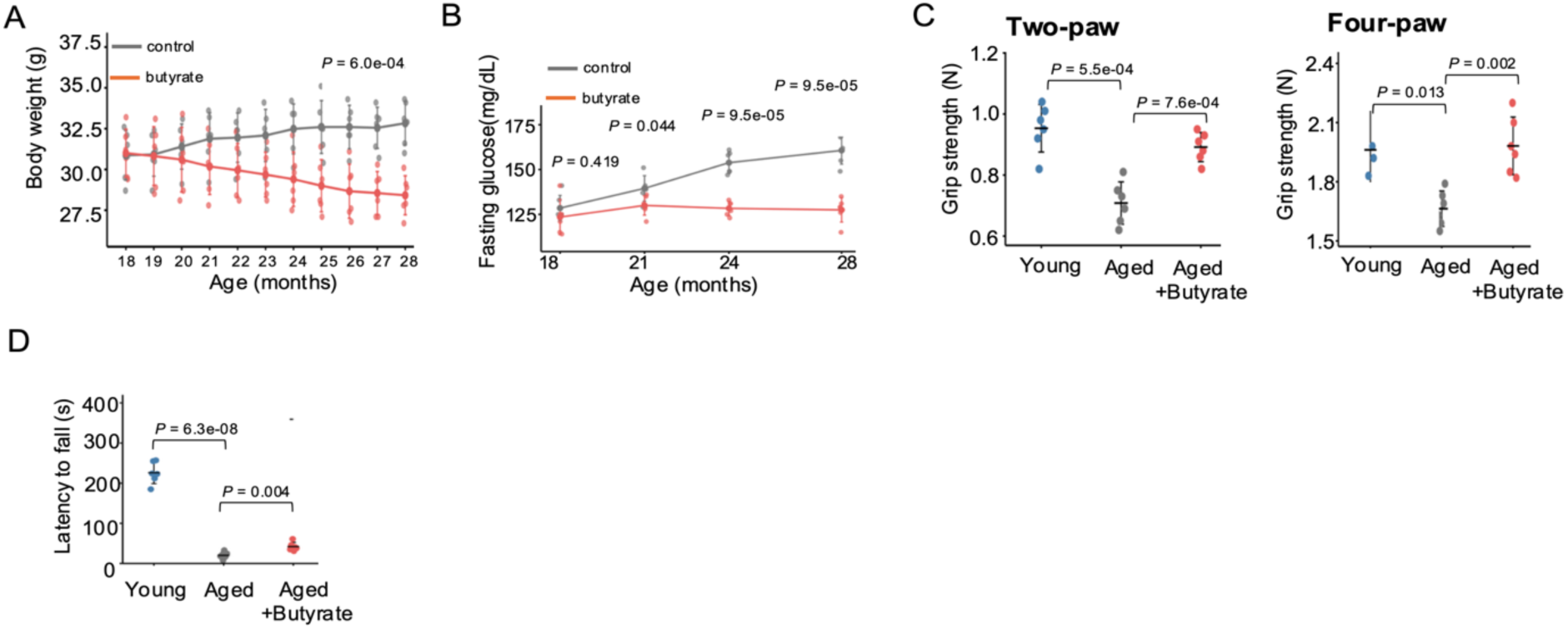
Late-life sodium butyrate supplementation improves metabolic and physical phenotypes in female mice. (A) Longitudinal body weight trajectories in female control and butyrate-treated mice from 18 to 28 months of age (n = 6 per group). The indicated P value represents the between-group comparison at 28 months. (B) Longitudinal fasting blood glucose concentrations measured at 18, 21, 24, and 28 months of age in female control and butyrate-treated mice (n = 6 per group). Indicated P values represent age-specific between-group comparisons. (C) Two-paw and four-paw grip strength in young, aged control, and aged butyrate-treated to fall during the final trial of an accelerating rotarod protocol in young, aged control, and aged butyrate-treated female mice (n = 6 per group). Young reference mice were 4 months old, and aged mice were 28 months old at physical-performance assessment female mice (n = 6 per group). Both measurements were obtained from the same animals. (D) Latency. Individual points represent biologically independent mice, and data are presented as mean ± SD. Longitudinal curves connect group means. Between-group comparisons were performed using two-sided, independent, equal-variance Student’s t-tests, followed by BH correction. Longitudinal P values represent comparisons at individual ages, rather than an overall treatment-by-time effect.

**Supplementary Figure 2.**
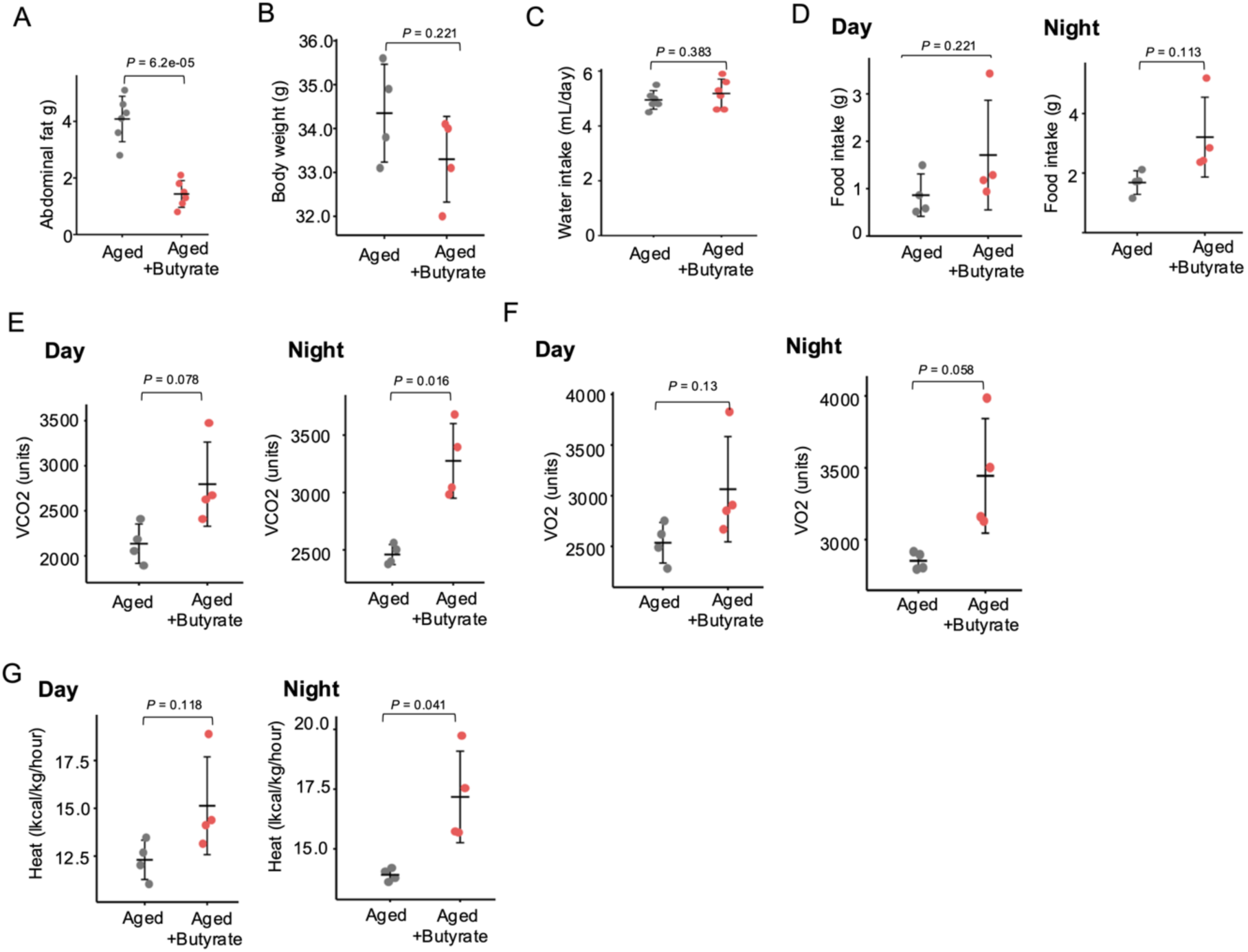
Late-life sodium butyrate supplementation reduces abdominal adiposity and alters whole-body respiratory metabolism in aged male mice. (A) Absolute combined epididymal and perirenal adipose tissue mass in aged male control and butyrate-treated mice, determined by dissection and weighing (n = 6 per group). (B) Body weight measured during the indirect-calorimetry assessment in aged male control and butyrate-treated mice (n = 4 per group). (C) Daily water intake in aged male control and butyrate-treated mice (n = 4 per group). (D) Food intake during the light (day) and dark (night) phases in aged male control and butyrate-treated mice (n = 4 per group). (E) Carbon dioxide production (VCO₂) during the light and dark phases in aged male control and butyrate-treated mice (n = 4 per group). (F) Oxygen consumption (VO₂) during the light and dark phases in aged male control and butyrate-treated mice (n = 4 per group). (G) Body-weight-normalized heat production during the light and dark phases in aged male control and butyrate-treated mice (n = 4 per group). Indirect calorimetry was performed in approximately 20-month-old male mice using a Comprehensive Lab Animal Monitoring System (CLAMS). Individual points represent biologically independent mice; summary marks and error bars indicate mean ± SD. Repeated calorimetry observations were summarized at the animal level before analysis. Two-sided, independent, equal-variance Student’s t-tests were used, with BH adjustment across the defined comparison families. Light-and dark-phase measurements were evaluated separately. Displayed P values were adjusted using the Benjamini–Hochberg adjustment. Heat production was normalized to body weight and expressed in kcal/kg/h.

**Supplementary Figure 3.**
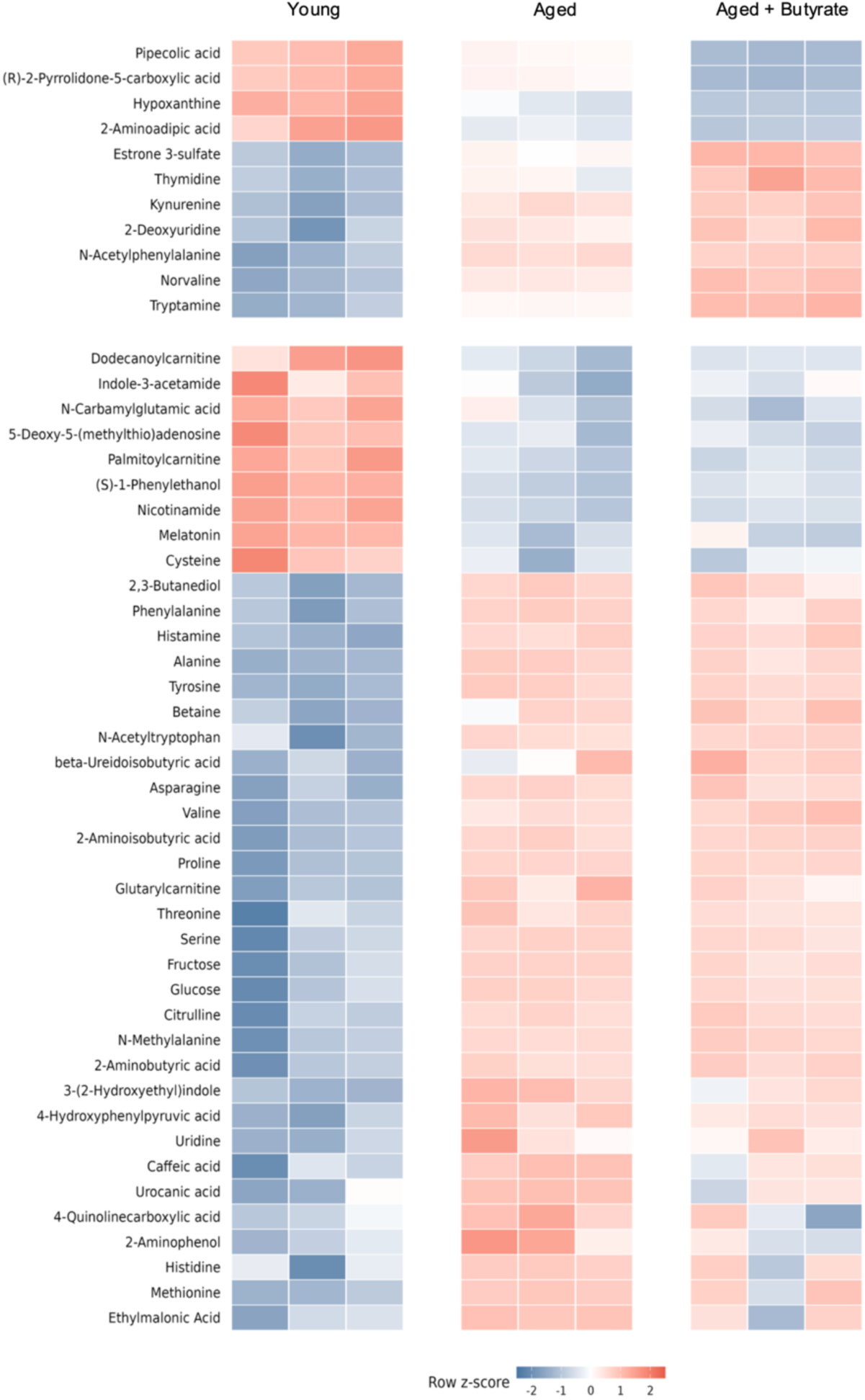
Additional circulating metabolite signatures associated with aging and late-life sodium butyrate supplementation. Related to Figure 2. Heatmap of selected plasma metabolites identified by broader metabolomic profiling in young (4-month-old), aged control (28-month-old), and aged sodium butyrate-treated (28-month-old) male C57BL/6J mice (n = 3 biologically independent mice per group). The upper portion highlights metabolites displaying distinct age-and treatment-associated abundance patterns, including pipecolic acid, (R)-2-pyrrolidone-5-carboxylic acid, hypoxanthine, 2-aminoadipic acid, estrone 3-sulfate, thymidine, kynurenine, 2-deoxyuridine, N-acetylphenylalanine, norvaline, and tryptamine. The middle and lower portions illustrate additional metabolites associated with amino acid metabolism, acylcarnitine metabolism, nucleotide metabolism, aromatic compound metabolism, and other biochemical processes, including metabolites whose age-associated changes persisted or differed in magnitude following butyrate supplementation. Each column represents an individual mouse, and each row represents a metabolite. Heatmap colors indicate row z scores calculated independently for each metabolite across the displayed samples, with red and blue denoting higher and lower relative abundance, respectively.

**Supplementary Figure 4.**
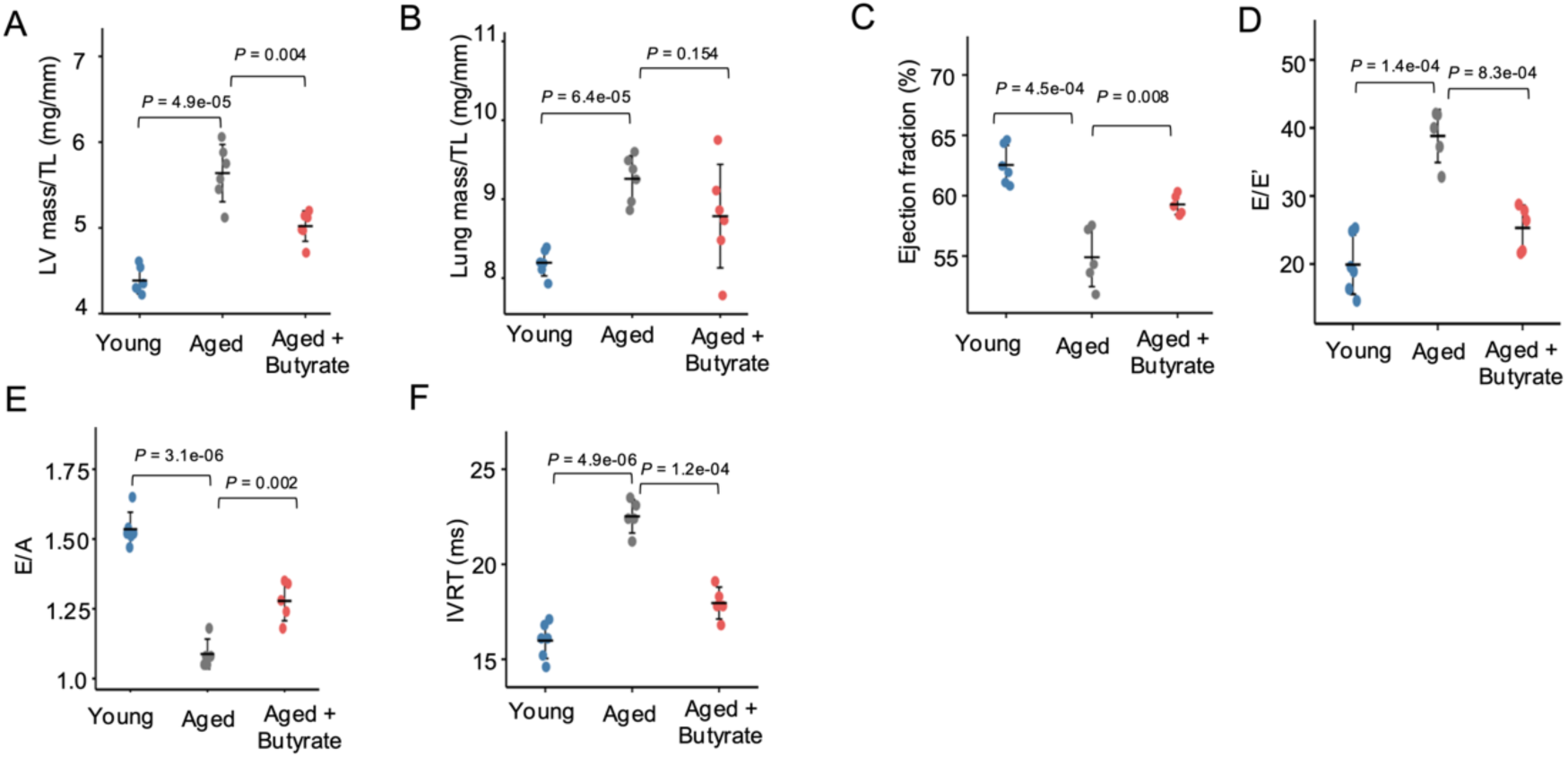
Late-life sodium butyrate supplementation attenuates age-associated cardiac remodeling and functional impairment in female mice. (A) LV mass normalized to TL, determined by weighing dissected LV tissue and expressed in mg/mm. (n = 6 biologically independent mice per group). (B) Lung mass normalized to tibial length (mg/mm). mice (n = 6 biologically independent mice per group). (C) LV ejection fraction (%) determined by echocardiography. (D) Ratio of E/E′ (E) Ratio of E/A (F) IVRT, expressed in milliseconds. Echocardiography: Young n=6, Aged n=5, Aged + butyrate n=5. Each point represents an individual mouse; horizontal summary marks and error bars indicate mean ± SD. Young reference mice were 4 months old, and aged animals were assessed at 28 months. Brackets indicate comparisons of young versus aged control mice and aged control versus aged butyrate-treated mice. Between-group comparisons were performed using two-sided, independent, equal-variance Student’s t-tests. Displayed P values were adjusted using the Benjamini–Hochberg adjustment. E, early transmitral filling velocity; A, late transmitral filling velocity; E′, early diastolic mitral annular velocity; LV, left ventricle; TL, tibial length; IVRT, isovolumic relaxation time.

**Supplementary Figure 5.**
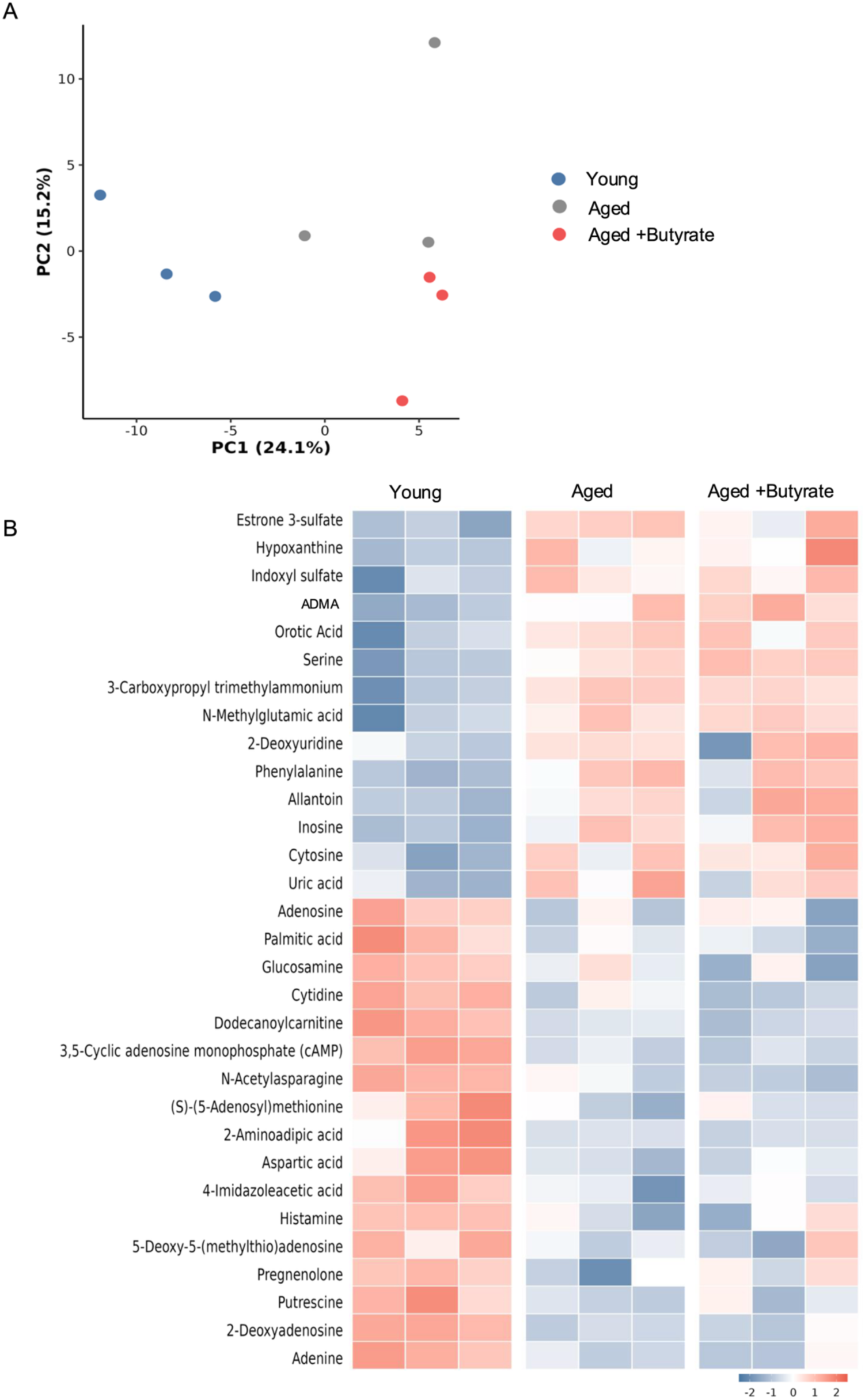
Additional myocardial metabolomic profiles associated with aging and late-life butyrate supplementation. LV metabolomic profiling was performed in young (4-month-old), aged control (28-month-old), and aged sodium butyrate-treated (28-month-old) male C57BL/6J mice (n = 3 biologically independent mice per group). (A) Principal component analysis (PCA) of LV metabolomic profiles. Each point represents an independent mouse. The first and second principal components explain 24.1% and 15.2% of total variance, respectively. (B) Heatmap of additional LV metabolites exhibiting age-and treatment-associated abundance patterns, including amino acid derivatives, nucleotide-related metabolites, acylcarnitines, organic acids, and other intermediary metabolic compounds. Each column represents an independent mouse, and each row represents a metabolite. Colors indicate row-standardized abundance, with red and blue corresponding to higher and lower relative abundance within each metabolite, respectively. The displayed metabolites illustrate both age-associated abundance differences that persist following treatment and selective treatment-associated changes.

**Supplementary Figure 6.**
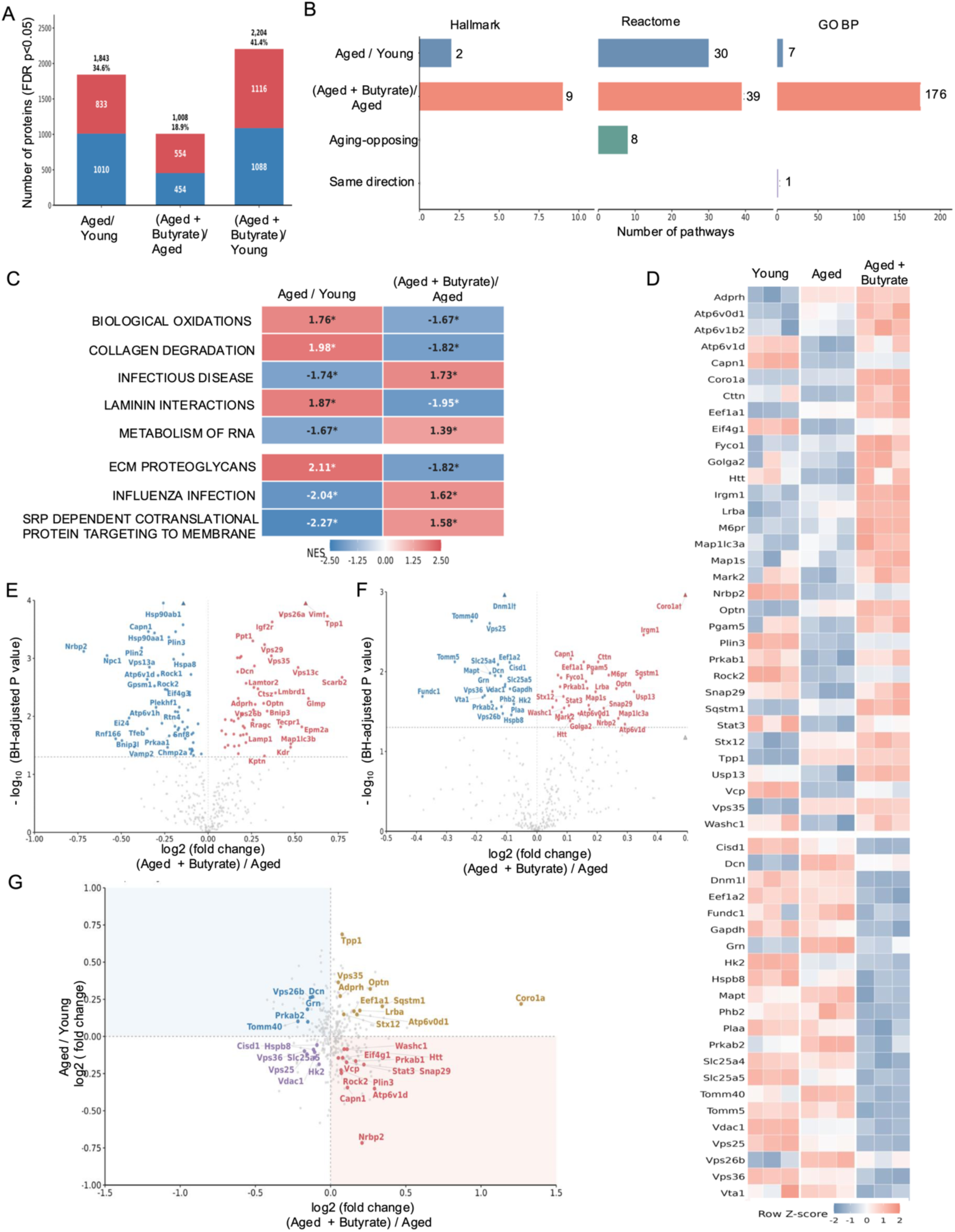
Proteomic remodeling and changes in autophagy-associated protein programs following late-life sodium butyrate supplementation. Left ventricular proteomics was performed in young (4-month-old), aged control (28-month-old), and aged sodium butyrate-treated (28-month-old) male C57BL/6J mice (n = 3 biologically independent mice per group). (A) Numbers of differentially abundant proteins identified in comparisons of aged control versus young, aged butyrate-treated versus aged control, and aged butyrate-treated versus young mice. Significantly altered proteins were defined using Benjamini– Hochberg-adjusted FDR < 0.05. Red and blue indicate proteins with higher and lower abundance, respectively, in the numerator group. Percentages above bars indicate the proportion of differentially abundant proteins relative to the analyzed protein set. (B) Numbers of significantly enriched pathways identified using Hallmark, Reactome, and Gene Ontology Biological Process (GO BP) gene-set collections in aging-and treatment-associated comparisons. Aging-opposing pathways exhibit statistically significant enrichment in opposite directions in aged control versus young and aged butyrate-treated versus aged control mice. Same-direction pathways exhibit significant enrichment in the same direction in both comparisons. (C) Normalized enrichment scores (NESs) for eight Reactome pathways showing significant enrichment in opposite directions between aging- and treatment-associated comparisons. Positive and negative values indicate enrichment in opposite directions, and asterisks denote pathways meeting the specified FDR threshold. (D) Row-standardized heatmap of selected proteins associated with autophagy, lysosomal function, vesicle trafficking, autophagosome formation, and related cellular quality-control processes. Each column represents an individual mouse. (E and F) Volcano plots highlighting selected autophagy-associated proteins in aged control versus young mice (E) and aged butyrate-treated versus aged control mice (F). The horizontal axis represents log₂ fold change, and the vertical axis represents −log₁₀(Benjamini–Hochberg-adjusted P value). Differentially abundant proteins were defined using an adjusted FDR < 0.05, with selected autophagy-associated proteins annotated. (G) Comparison of age-associated and treatment-associated log₂ fold changes for selected autophagy-associated proteins. The horizontal axis represents aged butyrate-treated versus aged control mice, and the vertical axis represents aged control versus young mice. Proteins in the upper-left and lower-right quadrants exhibit changes in opposite directions between aging and supplementation, whereas the remaining quadrants represent concordant directional changes. Protein differential-abundance P values were adjusted using the Benjamini–Hochberg procedure, with FDR < 0.05 used to identify differentially abundant proteins. Pathway enrichment significance was assessed using FDR correction across the relevant gene-set collection.

**Supplementary Figure 7.**
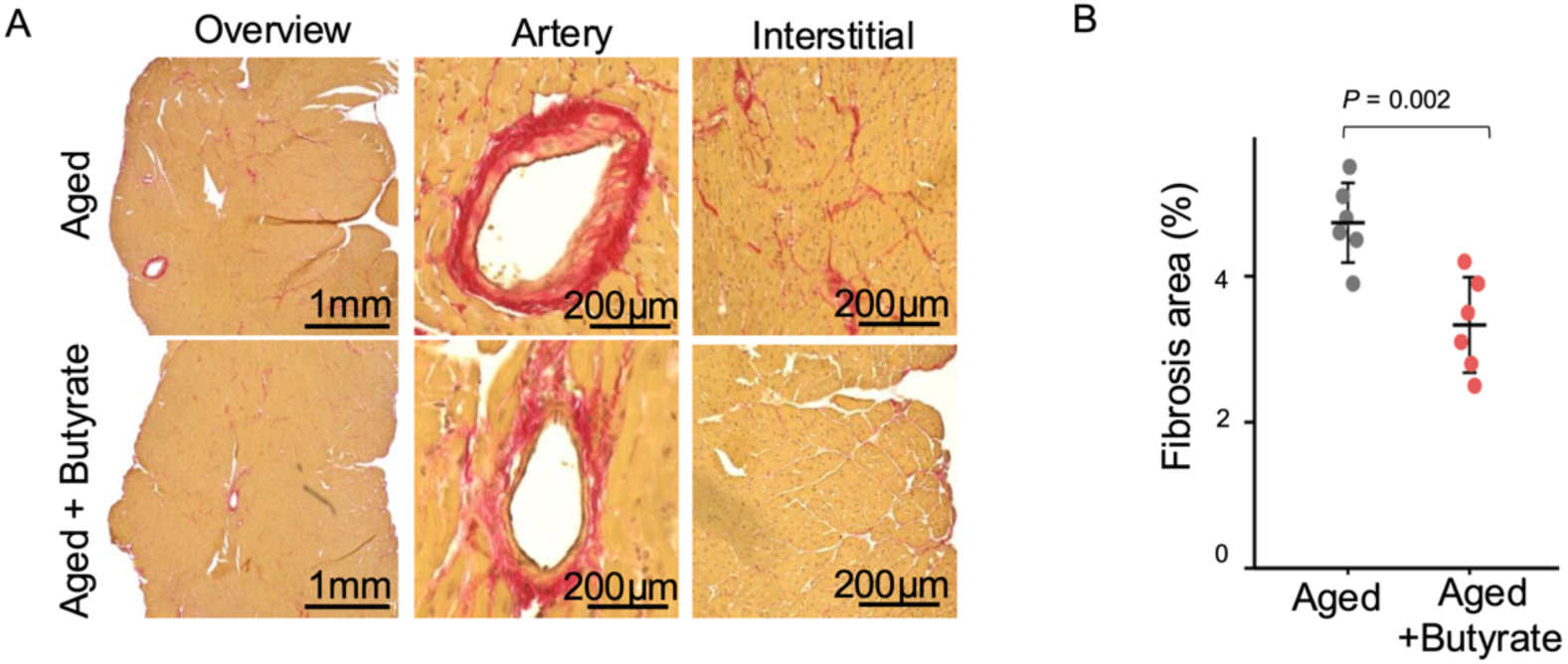
Late-life sodium butyrate supplementation attenuates myocardial fibrosis in aged mice. (A) Representative picrosirius red (PSR)-stained left ventricular sections from aged control and aged sodium butyrate-treated mice, showing whole-section overviews and higher-magnification images of perivascular and interstitial regions. Collagen-rich regions are visualized by red staining. Scale bars, 1 mm for whole-section images and 200 μm for higher-magnification images. (B) Quantification of collagen-positive fibrosis area, expressed as a percentage of the analyzed myocardial tissue area, in aged control and aged butyrate-treated mice (n = 6 biologically independent mice per group). Both perivascular and interstitial collagen-positive regions were included in the combined fibrosis measurement. Each point represents an individual mouse; horizontal summary marks and error bars indicate mean ± SD. Between-group comparisons were performed using two-sided, independent, equal-variance Student’s t-tests. The displayed P value (0.002) was adjusted using the Benjamini–Hochberg adjustment. PSR, picrosirius red; LV, left ventricle. Fibrosis area was quantified as the percentage of myocardial tissue area positive for collagen staining, using the image-analysis procedures.

**Supplementary Figure 8.**
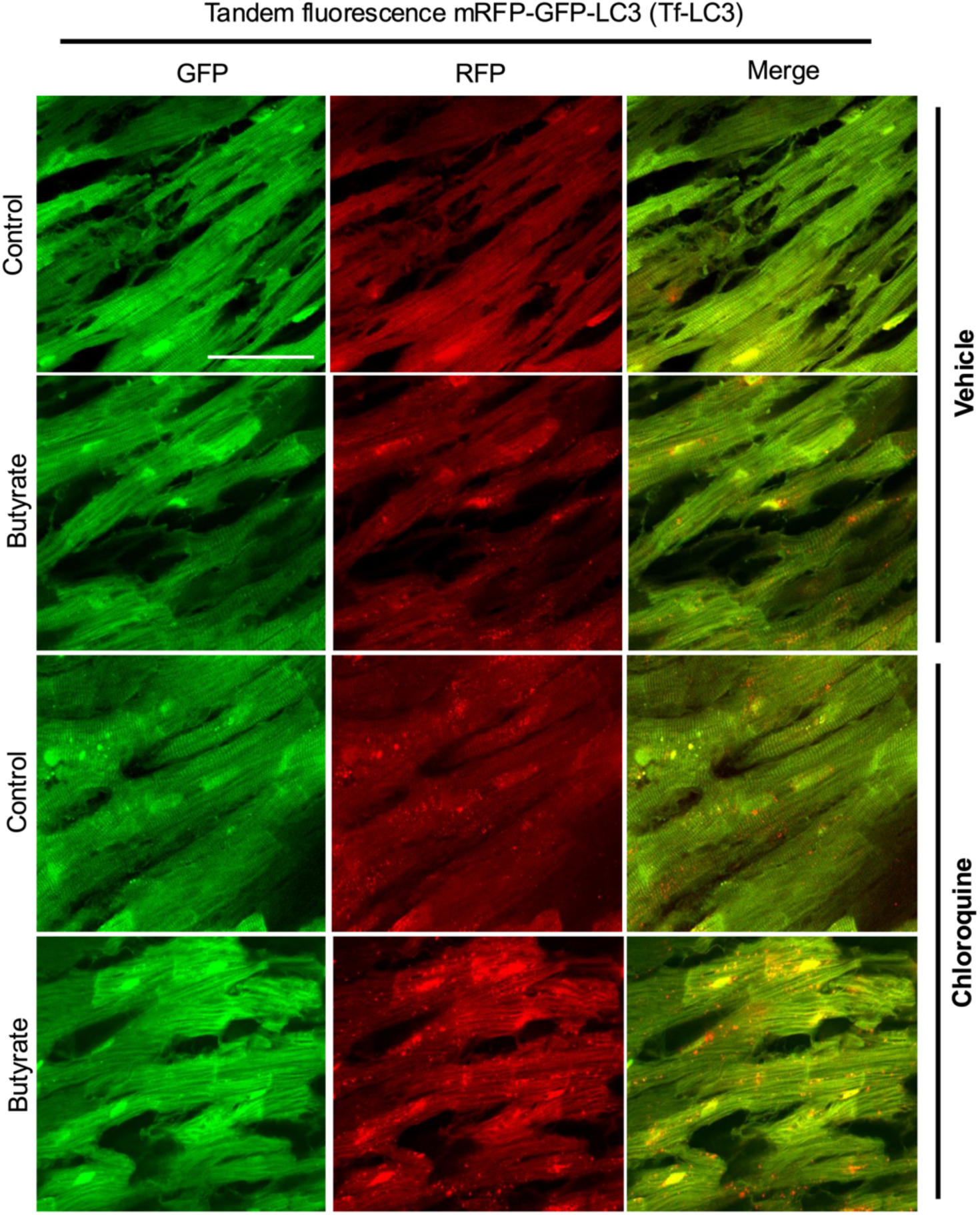
Additional tandem fluorescent LC3 images illustrating autophagic structures in control and butyrate-treated myocardium under vehicle and chloroquine conditions. Representative GFP, RFP, and merged images from the tandem fluorescent mRFP-GFP-LC3 (Tf-LC3) assay in control and butyrate-treated myocardium under vehicle and chloroquine conditions. GFP and RFP channels are shown separately together with the merged image to facilitate visualization of autophagosomes and autolysosomes. Autophagosomes appear as GFP-positive/RFP-positive puncta, whereas autolysosomes appear predominantly RFP-positive because GFP fluorescence is quenched in acidic lysosomal compartments. Chloroquine treatment was used to inhibit lysosomal degradation and aid interpretation of autophagic flux. Scale bar = 50µm.

